# Nutritional Anthelmintics: Chicory Reconfigures the Equine Holobiont Across Microbial, Parasitic, and Host Scales

**DOI:** 10.64898/2026.07.08.737212

**Authors:** Núria Mach, Sergio Méndez, Joshua Malsa, Juliette Auclair-Ronzaud, David Bars-Cortina, Marie-Amicie Sevillia-Boisferon, Gwendoline Pot, Maverick Monié--Ibanes, Hélène Henri, Oriane Chevalier, Corinne Régis, Camille Beaumelle, Antonio Velarde, Léa Lansade, Andrew R. Williams, Eric Richard, Glenn Yannic, Gilles Bourgoin, Géraldine Fleurance

## Abstract

Anthelmintic resistance in cyathostomins is escalating worldwide, threatening equine health and highlighting the need for sustainable, ecology-based parasite control strategies. Chicory (*Cichorium intybus, Puna II*) has emerged as a promising antiparasitic forage, yet its broader effects on the equine holobiont, parasites, microbiota, and host physiology remain poorly understood. We conducted a 32-day longitudinal grazing trial in young horses to assess how chicory affects parasitological outcomes, gut microbial ecology, nemabiome composition, behaviour, and host physiological and immune responses. Twenty-six naturally infected Anglo-Arabian horses were monitored weekly, with 13 grazing a chicory-based sward and 13 grazing a permanent pasture. Clinical parameters, body weight, and serum biochemistry remained stable across treatments, indicating that chicory was well tolerated. Immune profiles showed limited variation, although IL-10 increased in chicory-fed horses, suggesting subtle immune modulation. Behavioural observations revealed no signs of discomfort and indicated slightly enhanced social interactions in the chicory group.

Chicory grazing produced a marked reduction in cyathostomin egg excretion, accompanied by species-specific shifts in nemabiome composition. Several cyathostomin taxa, including *Cylicocyclus ashworthi, C. leptostomus,* and *C. nassatus*, declined in chicory-fed horses, whereas certain *Cylicostephanus* spp increased, indicating differential sensitivity rather than uniform suppression. Concomitantly, chicory induced profound ecological changes in the gut microbiota, including reduced alpha diversity, increased beta dispersion, and destabilised individual microbial trajectories. Several bacterial lineages, particularly *Oscillospiraceae, Clostridiaceae, Lachnospiraceae*, and Bacteroidales, were differentially enriched, reflecting a functional reorganisation of the intestinal ecosystem.

Together, these findings demonstrate that chicory reduces parasite fitness, reshapes nemabiome composition, and alters gut microbial ecology while maintaining host physiological stability. Chicory thus emerges as a promising ecological tool for parasite control, capable of modulating the equine holobiont in ways that complement and potentially reduce reliance on conventional anthelmintic strategies. However, because its effects on gut microbial ecology remain uncertain, and may include shifts resembling dysbiosis, future studies are needed to monitor microbial dynamics more closely and clarify the long-term ecological consequences of chicory grazing.

**Graphical abstract:** 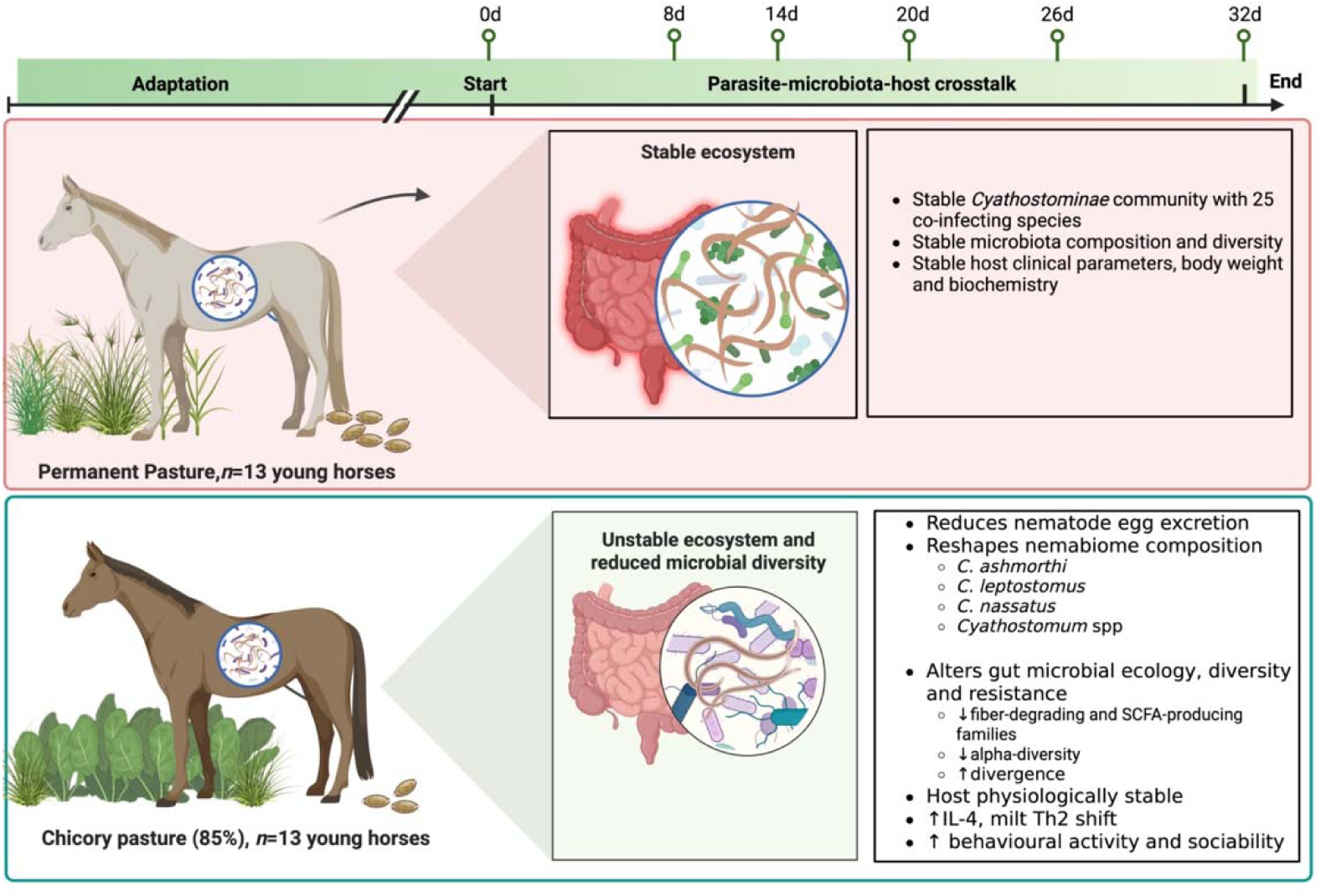

## INTRODUCTION

The holobiont concept considers mammals not as isolated organisms, but as integrated biological systems in which the host and its associated microbial and parasitic communities interact continuously to shape physiological function and homeostasis [1–3]. These interactions influence key processes such as nutrient metabolism, immune regulation, and resilience to environmental challenges [1–3]. In this light, the horse emerges as a dynamic holobiont in which tightly interconnected relationships among host tissues and microorganisms collectively determine health, welfare, and metabolic performance.

Within this ecological network, intestinal cyathostomin strongylids are ubiquitous and can impose substantial health and welfare burdens, including weight loss, colic, impaired immunity, and reduced performance [4–11]. Beyond their direct pathogenic effects, strongylid parasites play a key role in structuring gut ecology [12–15]. Helminths modulate mucosal immunity, influence mucus production, alter nutrient flows, and shape microbial communities through both direct and indirect mechanisms [16–18].

A growing body of work highlights the tripartite interplay between the gut microbiota, the nemabiome, and host immunity as a key determinant of equine health [14, 19–25]. Parasites can reshape microbial composition and metabolic outputs, while the microbiota influences parasite susceptibility, larval establishment, and immune tone [26]. Evidence for direct interactions between helminths and gut microbial species mainly concerns the reciprocal feedback between *Lactobacillaceae* and *Heligmosomoides polygyrus* [18] or *Trichuris muris* in mice [27], as well as the need for gut microbiota attachment to *T. muris* egg caps to trigger worm hatching [28].

In addition, parasite removal, whether through anthelmintics or dietary interventions, can also trigger ecological restructuring of the gut environment [14, 16, 26, 29, 30]. Studies in multiple host species show that parasite elimination often leads to shifts in microbial diversity, increased β dispersion, altered short□chain fatty acid production, and changes in inflammatory tone [26, 30, 31, 31–40]. These findings show that parasites are not only pathogens but also active structuring forces within the holobiont, shaping host physiology, microbial communities, and ecological interactions.

For decades, parasite control has relied almost exclusively on synthetic anthelmintics, but the rapid and widespread emergence of anthelmintic resistance now threatens the sustainability of equine health management [11, 41–45]. This crisis has stimulated growing interest in nutraceutical and plant based alternatives that can modulate parasite burden while supporting host physiology [46–51]. Plant secondary metabolites, including alkaloids, phenolics, and terpenes, may act through direct antiparasitic effects or by modulating host immunity, particularly by promoting Th2 responses involved in helminth control [49, 52, 53]. Although our recent work suggests that terpenes can exert *in vitro* antiparasitic and immunomodulatory effects and alter cyathostomin community structure, their *in vivo* efficacy in horses remains limited [49, 50], highlighting an incomplete understanding of their functional impact. This gap underscores the need to better characterise how plant-derived compounds operate within the complex holobiont.

Chicory (*Cichorium intybus, cultivar Puna II*) has emerged as a promising candidate for sustainable parasite control due to its rich content of sesquiterpene lactones, inulin, and polyphenols, compounds known for antiparasitic, anti□inflammatory, and prebiotic properties [54–56]. In ruminants, grazing on chicory at least 70% DM reduces gastrointestinal nematode burden [55, 57, 58]. Similarly, in horses, inclusion of 89% chicory in the diet reduces cyathostomin egg excretion and larval development [50].

However, the interactive effects of diet on the nemabiome, the gut microbiota, and host immunophysiology are mostly unknown. Understanding how a targeted nutritional intervention modulates this cross-kingdom network requires an integrative, multi-omics approach capable of capturing microbial, parasitic, behavioural, and physiological responses simultaneously. Building on this framework, the present study investigates whether a targeted plant-based intervention (chicory grazing) can modulate not only parasite burden but also the broader microbial and immunophysiological landscape. In this study, we conducted a longitudinal, multi-layered investigation combining faecal microbiota profiling (16S rRNA sequencing), nemabiome characterisation (ITS2 metabarcoding), faecal egg counts, behavioural assessments, and a comprehensive panel of host biomarkers. Our objective was to determine whether chicory grazing reshapes the equine holobiont and to identify the microbial, parasitic, immune, and physiological signatures associated with its antiparasitic effect. We hypothesised that chicory would (i) reduce strongylid burden and alter nemabiome composition; (ii) induce measurable shifts in the gut microbiota, including diversity and community structure; (iii) modulate host immune and metabolic responses, reflecting changes in gut ecology; and (iv) influence behavioural patterns, potentially reflecting broader changes in parasite burden and gut ecosystem functioning.

By integrating these datasets, we provide the first holistic demonstration of how chicory influences the equine holobiont and offer new perspectives for sustainable parasite control.

## RESULTS

### Experimental design and subjects

We monitored 26 naturally infected Anglo-Arabian young horses over a 32-day longitudinal grazing period to characterise their parasitological, microbial, physiological and behavioural dynamics. All horses had received no anthelmintics for 137 days prior to inclusion, corresponding to the interval since their last treatment and ensuring a stable, naturally acquired infection. At the start of the study, infections were dominated by small strongyles (*Cyathostominae*) as indicated by faecal egg counts (FEC; Additional Table 1).

**Table 1.**
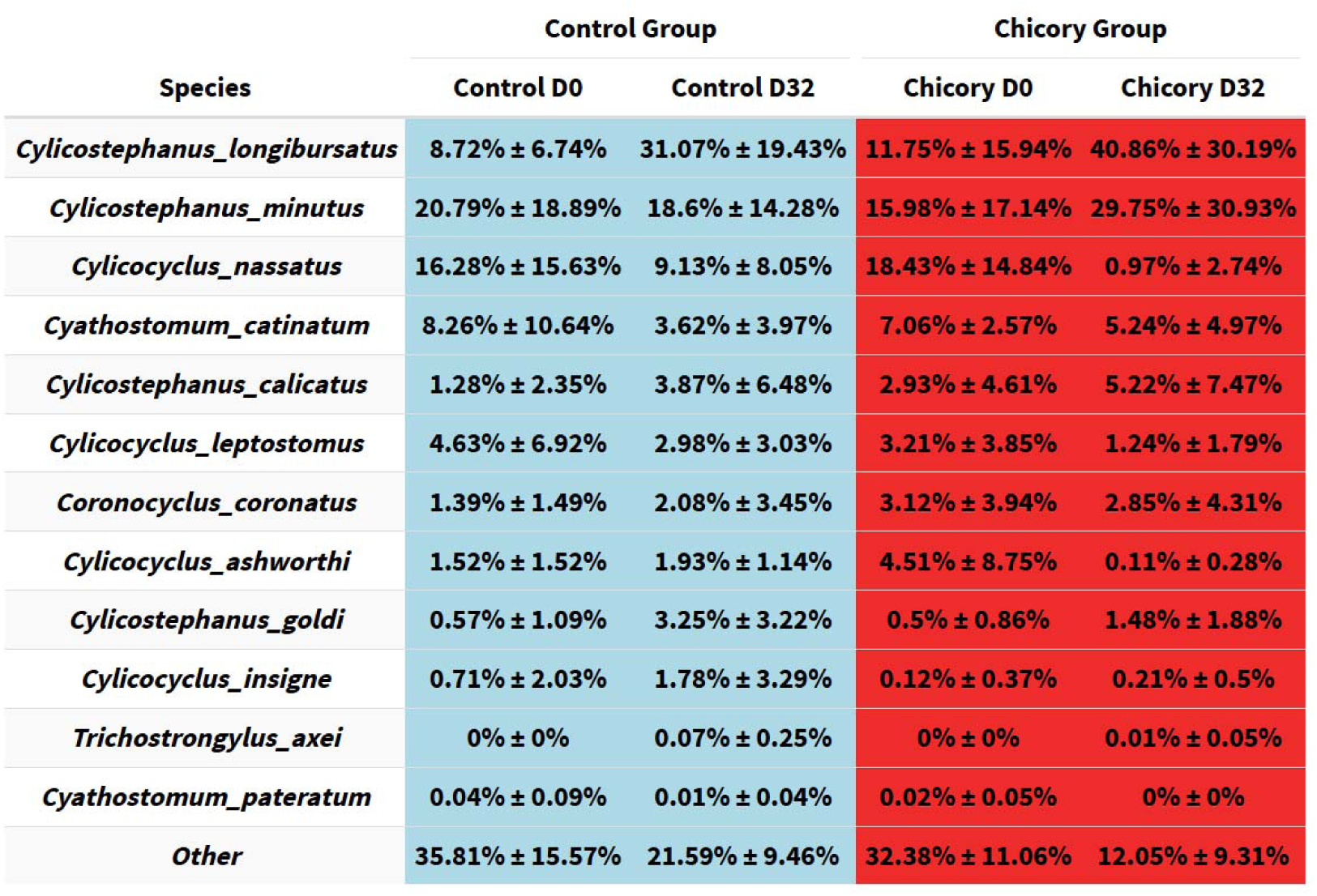
Comparison of the relative abundance (%) of the 12 nemabiome species identified at Day0 and Day32.

| Species | Control Group |  | Chicory Group |  |
| --- | --- | --- | --- | --- |
|  | Control D0 | Control D32 | Chicory D0 | Chicory D32 |
| *Cylicostephanus_longibursatus* | 8.72% ± 6.74% | 31.07% ± 19.43% | 11.75% ± 15.94% | 40.86% ± 30.19% |
| *Cylicostephanus_minutus* | 20.79% ± 18.89% | 18.6% ± 14.28% | 15.98% ± 17.14% | 29.75% ± 30.93% |
| *Cylicocyclus_nassatus* | 16.28% ± 15.63% | 9.13% ± 8.05% | 18.43% ± 14.84% | 0.97% ± 2.74% |
| *Cyathostomum_catinatum* | 8.26% ± 10.64% | 3.62% ± 3.97% | 7.06% ± 2.57% | 5.24% ± 4.97% |
| *Cylicostephanus_calicatus* | 1.28% ± 2.35% | 3.87% ± 6.48% | 2.93% ± 4.61% | 5.22% ± 7.47% |
| *Cylicocyclus_leptostomus* | 4.63% ± 6.92% | 2.98% ± 3.03% | 3.21% ± 3.85% | 1.24% ± 1.79% |
| *Coronocyclus_coronatus* | 1.39% ± 1.49% | 2.08% ± 3.45% | 3.12% ± 3.94% | 2.85% ± 4.31% |
| *Cylicocyclus_ashworthi* | 1.52% ± 1.52% | 1.93% ± 1.14% | 4.51% ± 8.75% | 0.11% ± 0.28% |
| *Cylicostephanus_goldi* | 0.57% ± 1.09% | 3.25% ± 3.22% | 0.5% ± 0.86% | 1.48% ± 1.88% |
| *Cylicocyclus_insigne* | 0.71% ± 2.03% | 1.78% ± 3.29% | 0.12% ± 0.37% | 0.21% ± 0.5% |
| *Trichostrongylus_axei* | 0% ± 0% | 0.07% ± 0.25% | 0% ± 0% | 0.01% ± 0.05% |
| *Cyathostomum_pateratum* | 0.04% ± 0.09% | 0.01% ± 0.04% | 0.02% ± 0.05% | 0% ± 0% |
| *Other* | 35.81% ± 15.57% | 21.59% ± 9.46% | 32.38% ± 11.06% | 12.05% ± 9.31% |

Horses were then allocated to either the chicory pasture or the control permanent pasture, with 13 horses per group. Allocation was balanced for age, sex, faecal egg counts (1020 ± 842 vs 1067 ± 837 FEC), and body weight (421.2 ± 33.1 vs 418.8 ± 29.7 kg) measured a few days before the start of the experiment, for chicory and control, respectively. The 32-day duration was intentionally kept shorter than the cyathostomin prepatent period to avoid reinfection and isolate the effects of the dietary intervention. Horses remained continuously on their assigned pastures, and sampling was performed at six predefined time points (D0, D8, D14, D20, D26, D32), covering parasitism outputs, behaviour, physiological parameters, and microbiota. This design enabled high-resolution characterisation of host–parasite–microbiome interactions under controlled yet natural grazing conditions (Figure 1AC).

**Figure 1.**
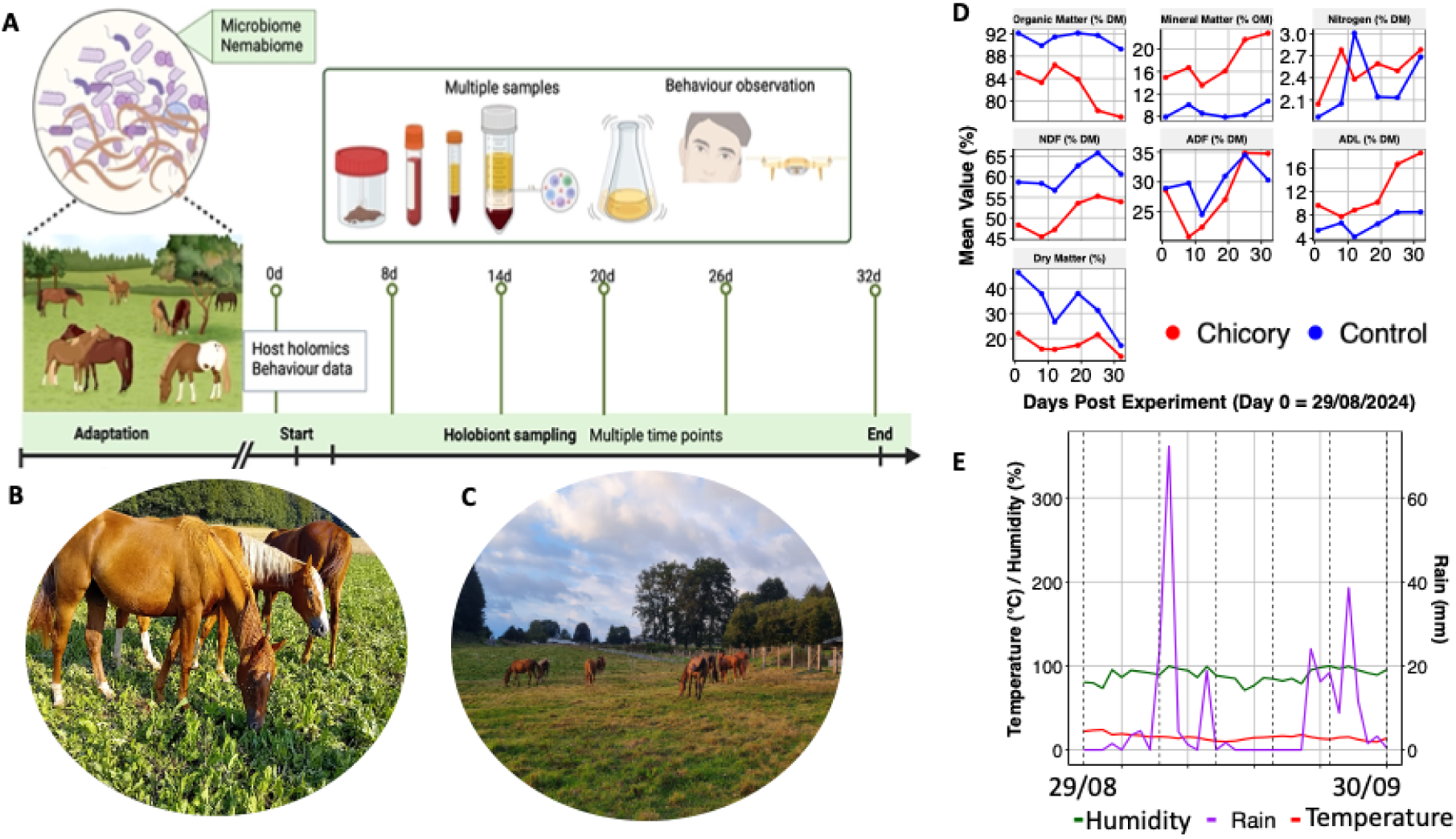
Experimental design, pasture conditions, and environmental context of the trial. **(A)** Schematic overview of the experimental workflow. Horses were turned out to pasture in April on different permanent grassland plots according to their age class and were subsequently regrouped onto a single plot one month prior to the start of the trial. During the experiment, horses from the two treatments (Chicory vs Control permanent pasture) were sampled at six predefined time points (D0, e.g., a few hours before entering the experimental plots; D8, D14, D20, D26, D32). At each time point, multiple sample types (faeces, blood, serum, and microbiota) were collected, together with behavioural observations, to characterize host–parasite–microbiome interactions under natural grazing conditions; **(B)** Horses grazing on chicory pasture, characterized by a dense sward and high leaf biomass; **(C)** Horses grazing on permanent grass pasture (control), showing a more heterogeneous vegetation structure; **(D)** Temporal evolution of chemical composition of plant biomass for chicor (red) and control (blue) treatments. Parameters include organic matter, mineral matter, nitrogen, neutral detergent fibre (NDF), acid detergent fibre (ADF), acid detergent lignin (ADL), and dry matter; and **(E)** Daily weather summary from 29 August to 30 September□2024 showing temperature (°C, red), humidity (%□green), and rainfall (mm, purple). Two rainfall peaks occurred around early and late September, but overall conditions remained warm and stable, supporting continuous grazing and consistent sampling.

### Pasture Structure and Quality Create Two Distinct Dietary Environments

Over the 32-day grazing period, the chemical composition of chicory and control pastures followed distinct, largely predictable trajectories, reflecting inherent botanical differences [59] rather than pronounced weather-driven fluctuations (Figure 1E). Organic matter remained consistently higher in the control pasture (>□88□%□DM), whereas chicory showed a gradual decline from ≈85□% to ≈75□%□DM (Wilcoxon test, *P* = 0.0163). This decrease was accompanied by a marked rise in mineral matter (up to ≈20□%□OM). Chicory exhibited a lower NDF content than permanent grassland. NDF, ADF, and ADL increased in both treatments over time, but the rise was steeper for NDF (Wilcoxon test, *P* = 0.0039) and ADL (lignin, Wilcoxon test, *P* = 0.0104) in chicory. Nitrogen content remained relatively stable across sampling days (2.1–3.0 % DM), with chicory consistently showing slightly higher values (Figure 1D).

The slightly shorter sward height observed in chicory plots (6.8□±□1.2□cm vs. 7.3□± 3.6□cm in controls) reflected intrinsic morphological differences rather than biomass depletion or rain-induced lodging.

### Faecal egg counts and Nemabiome dynamics during Chicory Grazing

#### Chicory grazing resulted in a clear reduction in faecal egg counts (FEC)

FEC were significantly lower in the chicory group at multiple time points (D8 and D18–D30; Wilcoxon test, *P* < 0.05), and this pattern remained consistent after normalisation to faecal dry matter (Figure 2A). In contrast to FEC, larval development was generally higher in the chicory group than in the control group, particularly at key time points. At D8, larval development wa markedly increased under chicory (65.5 ± 46.8%) compared to control (12.3 ± 3.9%; Welch’s t-test: *t*(11.2) = 3.93, *p* = 0.0023). A similar pattern was observed at D20, with greater development in the chicory group (29.9 ± 17.6%) than in the control (18.6 ± 4.9%; *t*(14.0) = 2.23, *p* = 0.0427). At D30, larval development remained elevated under chicory (28.9 ± 19.4%) compared to control (17.8 ± 5.2%), although this difference did not reach statistical significance (*p* = 0.0665). No significant differences were detected at D0 (*p* = 0.149), D14 (*p* = 0.429), or D26 (*p* = 0.605. This apparent discrepancy between reduced FEC and increased larval development under chicory coincided with marked changes in the intestinal physicochemical environment. Faecal redox potential was significantly reduced from D14 onwards (*P* < 0.0001), indicating a shift toward a more reducing and fermentative gut environment (Figure 2B). Faecal pH remained within the physiological range but showed a transient increase in the chicory group between D12 and D18 (Figure 2C).

**Figure 2.**
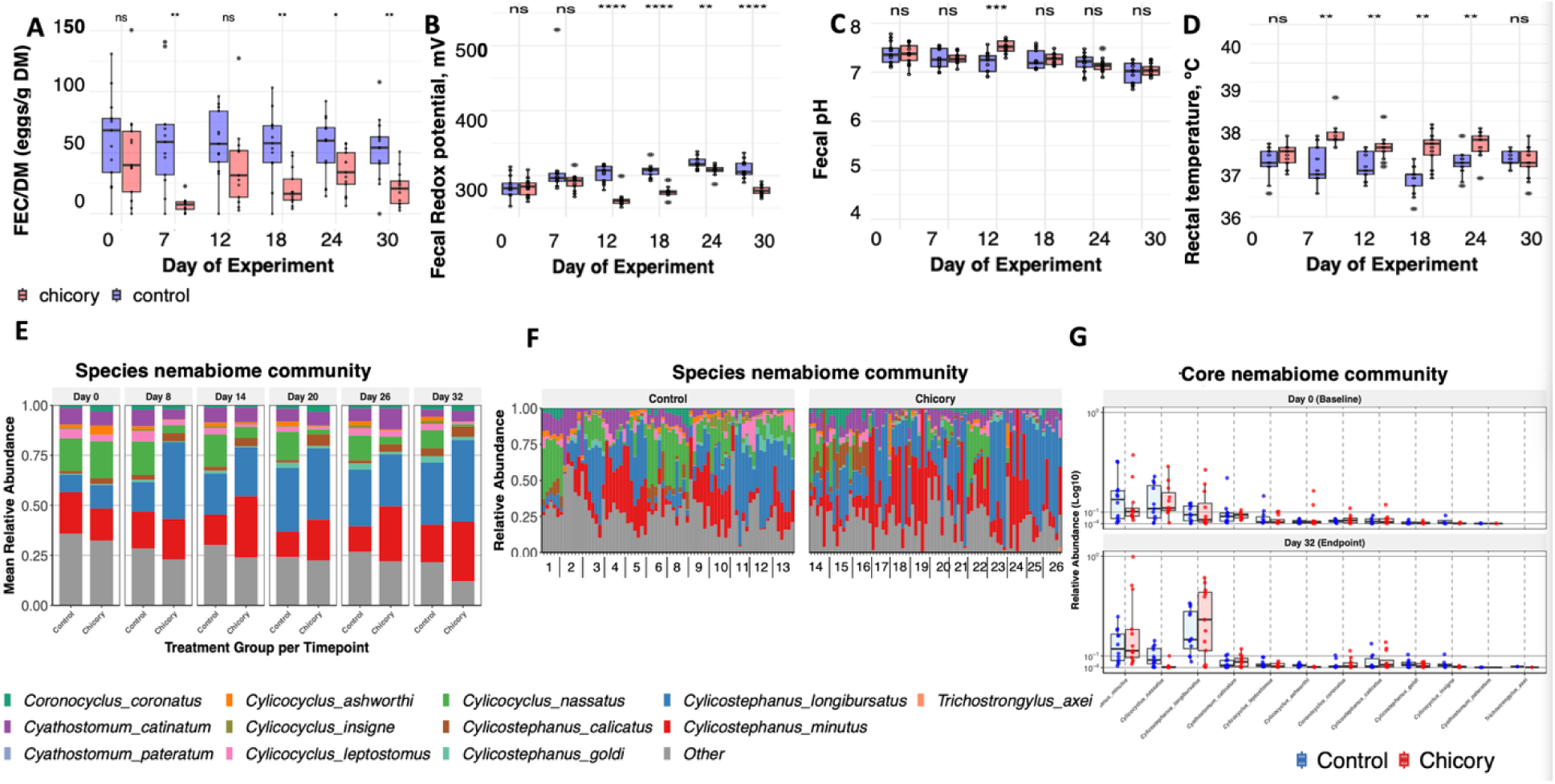
Effects of chicory grazing on parasite burden, gut environment, and nemabiome composition. **(A)**□Boxplots show faecal egg counts (FEC, eggs/g faecal dry matter) across timepoints for chicory-fed (red) an control (blue) horses. Each box represents the interquartile range, the horizontal line indicates the median, and the whiskers denote the spread of the data; **(B)**□Boxplots illustrate faecal redox potential (mV) over time; **(C)**□Boxplots display faecal pH dynamics; **(D)**□Boxplots show rectal temperature (°C) across timepoints; **(E)**□Stacked bar plots show mean relative abundance of cyathostomin species across timepoints, revealing a stable core nemabiome; **(F)** Individual-level nemabiome composition across all horses in the control and chicory groups. Each vertical bar represents one horse. Colours correspond to species identities. Others include unassigned ASV at the species level; and **(G)** Boxplots represent the core cyathostomin nemabiome in both groups, showing the relative abundance of the dominant species shared across individuals and time points. Each box illustrates the interquartile range, the median is shown by the central line, and whiskers indicate data dispersion.

#### Nemabiome Composition and Core Community Structure

To assess whether the marked reduction in cyathostomin egg excretion was accompanied by changes in parasite community composition, the nemabiome was characterised using ITS-2 rDNA metabarcoding. From ≈20.6□million sequence (138,227.9□±□83,442.3□reads□per□sample), 6,649□ASVs were identified, of which 3,637□(54.7%) were confidently assigned to the species level. Aggregation yielded 37□species-level taxa, including 24□co-infecting *Cyathostominae* species shared between treatments (Additional Table 2). After refinement, 12 cyathostomin taxa were described at the species level, accounting for 65–79% of all reads, forming a highly dominant and biologically coherent core community (Figure□2E; Table□1). As in previous equine nemabiome studies [5, 51, 60, 61], baseline communities were dominated by *Cylicostephanus longibursatus*, *C. minutus*, and *Cylicocyclus nassatus*, each contributing >10% of total reads (Figure 2E), with additional core taxa such as *Cyathostomum catinatum*, *C. leptostomus*, *C. ashworthi*, *Coronocyclus coronatus* and *Cylicostephanus calicatus* present at lower but consistent relative levels (Figure 2G).

Interestingly, the species-level relative composition plots revealed marked inter-individual variability (Figure 2F), with horses displaying highly distinct cyathostomin profiles throughout the grazing period (Additional Table 2). Even within the same treatment group, the relative abundances of dominant taxa such as *Cylicostephanus longibursatus*, *C. minutus*, and *Cylicocyclus nassatu*s varied widely between individuals, producing idiosyncratic temporal trajectories rather than a uniform group-level pattern.

#### Nemabiome diversity and Ecological Stability

The nemabiome displayed remarkable ecological stability. Alpha diversity (Shannon index) remained constant across time and treatments (≈□2.0–4.0), showing that chicory grazing did not alter overall species richness or evenness. This stability in diversity was mirrored in beta-diversity patterns, where PCoA showed substantial overlap among treatments across all sampling days (Figure 3A-B). ANOSIM further confirmed low inter-group structural separation (R²□<□0.11□at all timepoints; □adj *P*□>□0.05, except D8□adj□*P*□=□0.030), indicating that chicory-fed and control horses maintained largely overlapping community structures throughout the trial (Additional Table 3). Distance-to-centroid analyses confirmed that both groups maintained tight, stable clustering throughout the trial, with no evidence of increased dispersion or stochastic drift (lmer, *P* > 0.05; Figure 3C). Complementary, measures of within-group consistency reinforced this interpretation: inter-individual variation (internal Bray–Curtis distances) remained low and unchanged from D0 to D32 (lmer, *P* > 0.05; Figure 3D).

**Figure 3.**
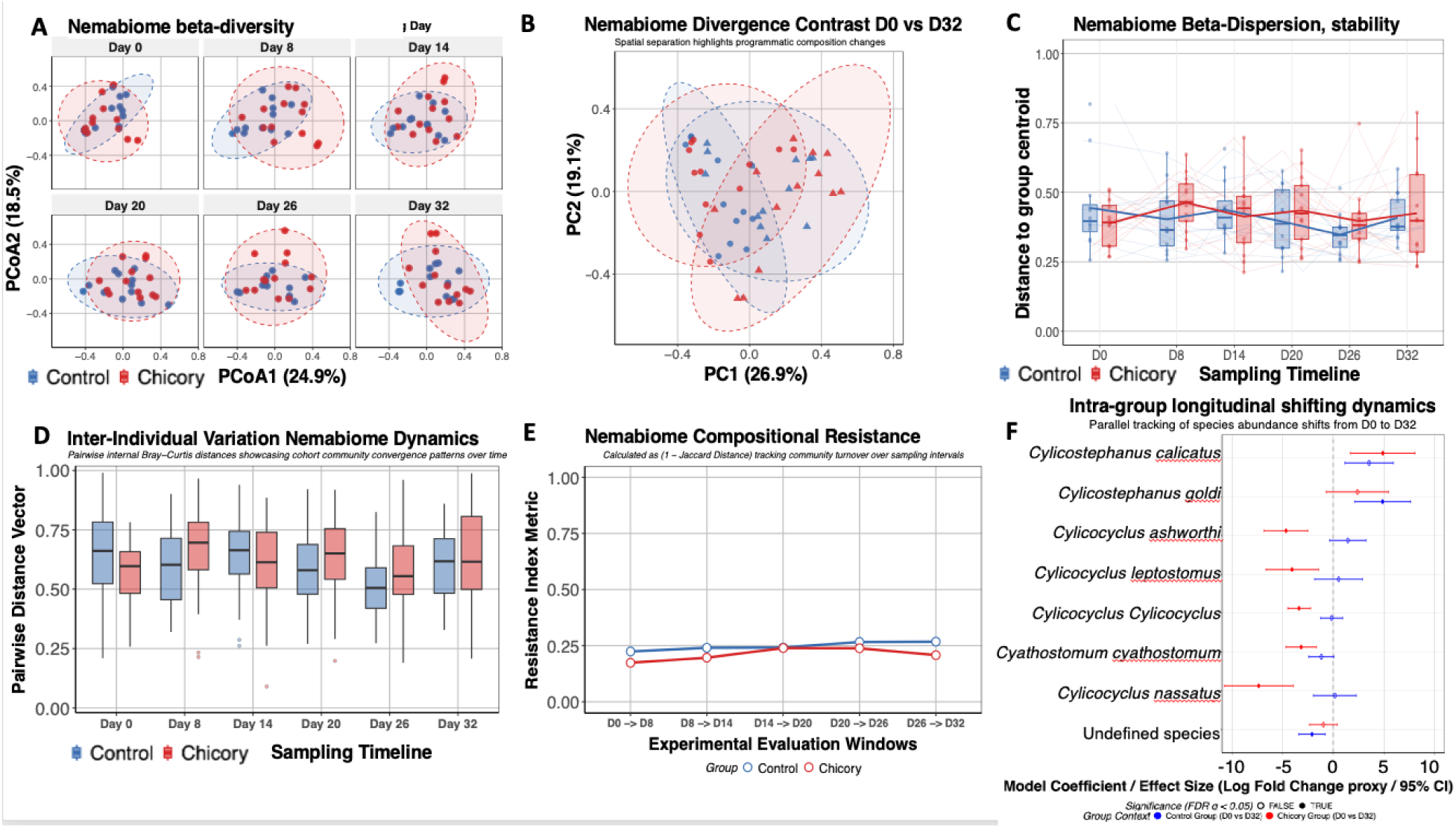
Nemabiome community dynamics under chicory grazing. **(A)** Principal coordinate analysis (PCoA) of Bray–Curtis distances illustrating temporal shifts in nemabiome β-diversity between control (blue) and chicory (red) groups across all sampling days. Ellipses represent 95□% confidence intervals for group clustering; **(B)** PCoA contrasting Day□0 and Day□32 samples, highlighting no spatial separation and compositional divergence between groups; **(C)** β-dispersion analysis showing temporal changes in community stability, expressed as distance to group centroid. Increasing dispersion indicates reduce structural homogeneity over time; **(D)** Inter-individual variation in nemabiome composition, represented by pairwise internal Bray–Curtis distances. Boxplots depict cohort-level convergence or divergence patterns throughout the experiment; **(E)** Nemabiome Compositional Resistance: the blue line with circular markers represents the control group, while the red line with square markers represents the chicory-treated group. These two colored lines track the resistance index. The resistance index reflects how stable the nematode communit composition remains over time, with higher values indicating greater stability; and **(F)** Results of a MaAsLin2 longitudinal model, which estimates how the relative abundance of each cyathostomin species changes within each treatment group over time. The model uses Day 0 as the reference, so each coefficient represents the direction an magnitude of change from baseline to Day 32. Positive coefficients → the species increased in relative abundance over the trial; negative coefficients → the species decreased relative to baseline. Error bars represent the 95% confidence interval around each estimate. This approach allows detection of true temporal trends within eac group, independent of between-group differences. ns□=□not significant; *□P□<□0.05; **□P□<□0.01; ***□P□<□0.001; ****□P□<□0.0001.

However, PerMANOVA analyses demonstrated significant divergence between chicory and control nemabiomes after grazing onset: no difference at baseline (Day□0,□R²□=□0.029, adj *P* =□0.580), but clear separation from Day□8 onward (R²□=□0.258–0.408; adj□*P*□≤□0.005). The strongest differentiation occurred at Day□20 (R²□=□0.408,□adj *P*□=□0.002). Finally, compositional resistance (1□–□Jaccard distance) demonstrated high temporal persistence in both groups, with resistance values of ≈□0.25 (Figure 3E).

#### Nemabiome species-Specific Restructuring under Chicory Grazing

Having characterised the overall community structure through alpha and beta diversity analyses, we next examined species-level relative patterns to determine which taxa were specifically driving these ecological shifts. As grazing progressed, the relative composition of parasite species trajectories diverged markedly between treatments. In the control group, the core nemabiome remained largely stable, with only a gradual increase in *Cylicostephanus longibursatus* relative abundance (from ≈10% at Day□0 to ≈31% at Day□32; adj *P* = 0.00158). In contrast, the chicory group underwent a much more pronounced restructuring in the core microbiome: *C.*□*nassatus* declined to near-background levels (<□4%; adj *P* <□0.05), indicating a strong treatment-driven suppression of these taxa (Figure 2E-F). This pattern was further clarified by MaAsLin2 longitudinal modelling, which revealed extensive species-specific shifts in the chicory group between Day□0 and Day□32 (Figure 3F). One species, *Cylicostephanus calicatus*, showed significant positive coefficients, reflecting increased relative abundance over time. In contrast, three species exhibited significant declines in relative composition under chicory grazing: *Cylicocyclus ashworthi*, *C.*□*leptostomus*, and *C.*□*nassatus* (all adj *P* <□0.05; Additional Table 4).

### Sustained Metabolic and Immune Host Stability Under Chicory Grazing Despite Changes in Parasite Burden

After characterising the rich and diverse cyathostomin community colonising the horses’ intestines, we next assessed whether such parasite loads, or their dietary treatment-driven shifts, were associated with any impact on host health or behaviour. To ensure that animals remained clinically stable, we monitored digestive symptoms, body weight, rectal temperature, metabolic markers, immune parameters and behaviour throughout the trial.

Across the experimental period, none of the animals exhibited gastrointestinal disorders or any clinical symptomatology, and chicory grazing maintained a remarkably stable metabolic and immune profile, with only mild and transient physiological adjustments. Specifically, body weight remained constant across treatments (430□±□25□kg vs□435□±□27□kg for chicory and control groups, respectively) while rectal temperature showed a slight but physiologically normal increase in chicory-fed horses (*P*□<□0.01), coinciding with small early rises in total plasma proteins (*P*□<□0.05–0.001 between D0 and D18; additional Table 1). Core metabolic markers, including bile acids (3.2□±□0.9 vs 3.0□±□1.0□µmol/L), β-hydroxybutyrate (0.26□±□0.05 vs 0.25□±□0.06□mmol/L), and non-esterified fatty acids (NEFA), remained stable. NEFAS exhibited only transient fluctuations, with slightly lower values in the chicory group at the beginning and mid-experiment (*P*□<□0.05 on D0 and D18), but no consistent differences thereafter (0.18□±□ 0.09□mmol/L vs□0.22□±□0.11□mmol/L). Protein fractions supported this metabolic stability: urea concentrations remained stable across all sampling points (0.43□±□0.07□g/L for chicory vs□0.46□±□0.08□g/L for controls). Albumin also remained unchanged (24.5□±□2.1 vs 25.0□±□2.3□g/L), while minor shifts were recorded in α□-globulins (decrease at mid-experiment, *P*□<□0.01) and α -globulins (increase later, *P*□<□0.001). When expressed as percentages, albumin and globulin proportions followed similar patterns, with albumin accounting for approximately 40□±□4% of total plasma proteins and globulins for 60□±□5%, maintaining a balanced albumin-to-globulin ratio across treatments.

Immune indicators further confirmed homeostasis. Total plasma proteins were slightly higher in chicory-fed horses during the early phase of the trial (*P*□<□0.05–0.001 between D0 and D18). SAA stayed low with only a small rise at D30 in both groups, and leukocyte profiles, including neutrophils (≈45–55%), eosinophils (≈4–6%), basophils (≈0.7□±□0.3%), monocytes (≈4–6%), and lymphocytes (≈40–50%), remained within physiological ranges with no treatment effects. Cytokine patterns aligned with this equilibrium. Cytokine responses generally reflected a stable immune state, with TNF-α remaining consistent across time points. Most Th2 cytokines showed limited variation; however, IL-4 demonstrated a significant increase in the chicory group at D20 and D32. In contrast, IL-5 and IL-13 did not exhibit meaningful changes. IL-10 displayed a transient mid-experiment elevation (P□<□0.01), suggesting a temporary enhancement of regulatory activity rather than a sustained inflammatory response (Additional Table 1).

### Chicory behavioural responses: increased engagement and reduced inactivity

We then considered whether either the parasitic load or the chicory diet itself could influence how animals interacted with each other and with their environment. Behaviour often reveals subtle changes in comfort and motivation, long before clinical or metabolic alterations appear [62, 63]. For this reason, we complemented our physiological assessments with a detailed evaluation of feeding patterns, foraging choices, locomotor activity, social interactions, and potential signs of discomfort, allowing us to determine whether chicory grazing shaped not only parasite dynamics but also the horses’ day-to-day behavioural responses.

Eating behaviour recorded with Ethosys® collars showed no consistent treatment effect. Across all monitoring days, total daily eating time was similar between groups (Chicory: 17□h□44□min; Control: 18□h□05□min; Wilcoxon: NS). However, scan-sampling during 7□h of direct observation revealed some differences (Figure 4A). On average, chicory-fed horses spent slightly but significantly higher proportion of time feeding (62.5% ± 8.1 vs. 60.3% ± 6.9, respectively, Wilcoxon *P* =□1.35□×□10□¹□, Figure□4A). Foraging behaviour diverged clearly between treatments: horses in the control group primarily grazed grasses, which constituted 79.2□±□6.6% of the bites consumed. Conversely, horses in the chicory group mainly selected chicory, with 77.2□±□14.4% of bites dominated by chicory and 80.3□±□12.2% containing chicory. Among the other plant species present in the plots with reported anthelmintic properties in ruminants, plantain (*Plantago lanceolata*) was the only species consumed by horses in both treatments. However, its contribution remained limited, accounting for 6.4□±□2.9% of bites in the control group and 1.8□±□2.0% in the chicory group (Wilcoxon test, *P*□<□0.0001).

**Figure 4.**
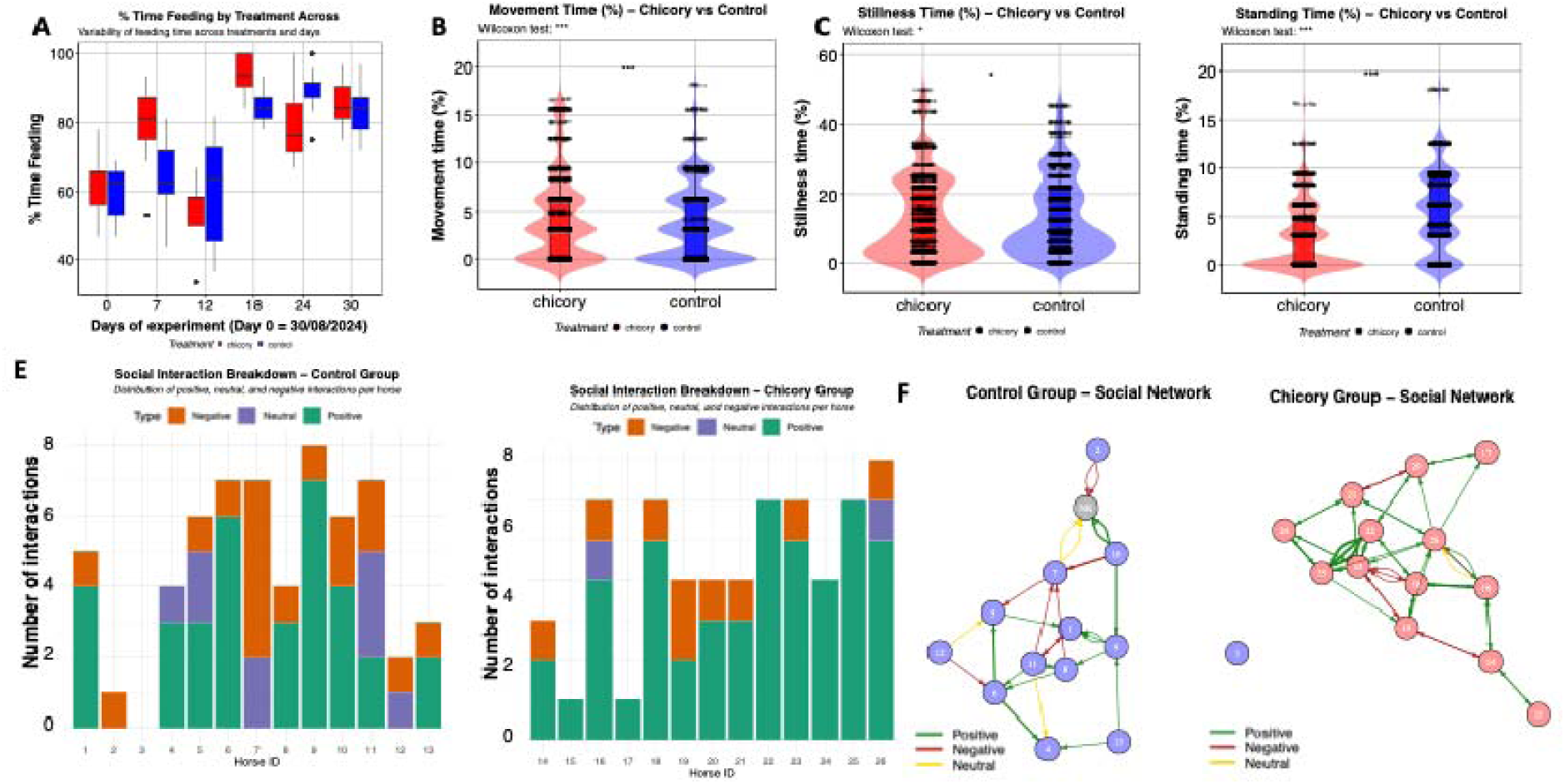
Behavioural and social patterns of horses grazing chicory or control pastures across the experimental period. Top panels: **(A)**□Percentage of time feeding (boxplots), **(B)**□time in movement, **(C)** Stillness□time t, and **(D)**□time standing, recorded by scan sampling. Red□=□chicory; blue□=□control; asterisks indicate significant differences between treatments. Bottom panels: (**E–F**)□Distribution of positive, neutral, and negative social interactions per horse in control and chicory groups; (**G–H**)□social network diagrams showing the density and valence of interactions (green□=□positive, yellow□=□neutral, red□=□negative).

Locomotor activity followed the same pattern. Chicory-fed horses spent a little more time moving (4.31% vs 3.68%; *P* =□0.00016) and a bit less stillness time (12.73% vs 13.07%; *P* =□0.0297; Figures□4B–C). Standing behaviour showed similar trends, with chicory horses standing slightly less overall compared to control counterparts (3.24% vs 5.44%; *P* =□0.0426, Figure□4D). Social interactions showed substantial interindividual variability, yet a qualitative pattern emerged when examining interaction valence and the shape of social networks (Figure 4E-F). Horses grazing chicory engaged in a higher proportion of positive interactions, such as affiliative contact, mutual sniffing, and close-proximity grazing, compared with controls, whose interactions included a slightly higher proportion of neutral or low-intensity agonistic behaviours. Although these differences were not statistically significant, the structure of the social networks provides additional insight: the chicory group displayed a denser, more evenly connected network, with more reciprocal links and fewer isolated individuals, suggesting a more cohesive and socially integrated group.

Lastly, indicators of potential discomfort during grazing remained low in both groups, with no difference in the proportion of grazing time spent with ears held back (Control: 0.7□±□1.2%; Chicory: 1.0□±□1.3%), suggesting that neither pasture type induced overt negative affective states.

### Microbiota composition, diversity, and ecological stability

Having shown that chicory grazing both reduced parasite burden and reshaped the underlying cyathostomin community, we next asked whether these shifts, together with the direct effects of chicory itself, also influenced the gut microbial ecosystem. We therefore characterised the faecal microbiota structure using 16S rRNA gene sequencing, quantified alpha and beta diversity, and assessed the ecological stability of microbial communities over time.

#### Baseline microbial taxonomic and core microbiome structure

After quality filtering, denoising, and chimaera removal, the dataset contained ∼7.7□×□10□high-quality 16S rRNA sequences (50,385.7□±□37,332.1 per sample) and 11,007 ASVs, which were subsequently collapsed into taxonomic lineages because many distinct ASVs corresponded to the same biological taxon, resulting in a reduced and biologically meaningful set of ∼433 unique lineages (Additional Table 5). At baseline, the faecal microbiota of horses was dominated by two major phyla: Bacillota (≈□44%) and Bacteroidota (≈□38%), together accounting for more than 80% of the total community. Minor contributors included Fibrobacterota (≈□2–3%), while Actinomycetota, Verrucomicrobiota, and other low-abundance phyla collectively represented less than 5% (Figure 5A). At the family level, the community was largely composed of key anaerobic fermenters, including *Prevotellaceae* (≈□14–17%), *Lachnospiraceae* (≈□15–17%), and *Oscillospiraceae* (≈□7–9%), followed by *Rikenellaceae* and *Christensenellaceae*, each contributing around 5–8%.

**Figure 5.**
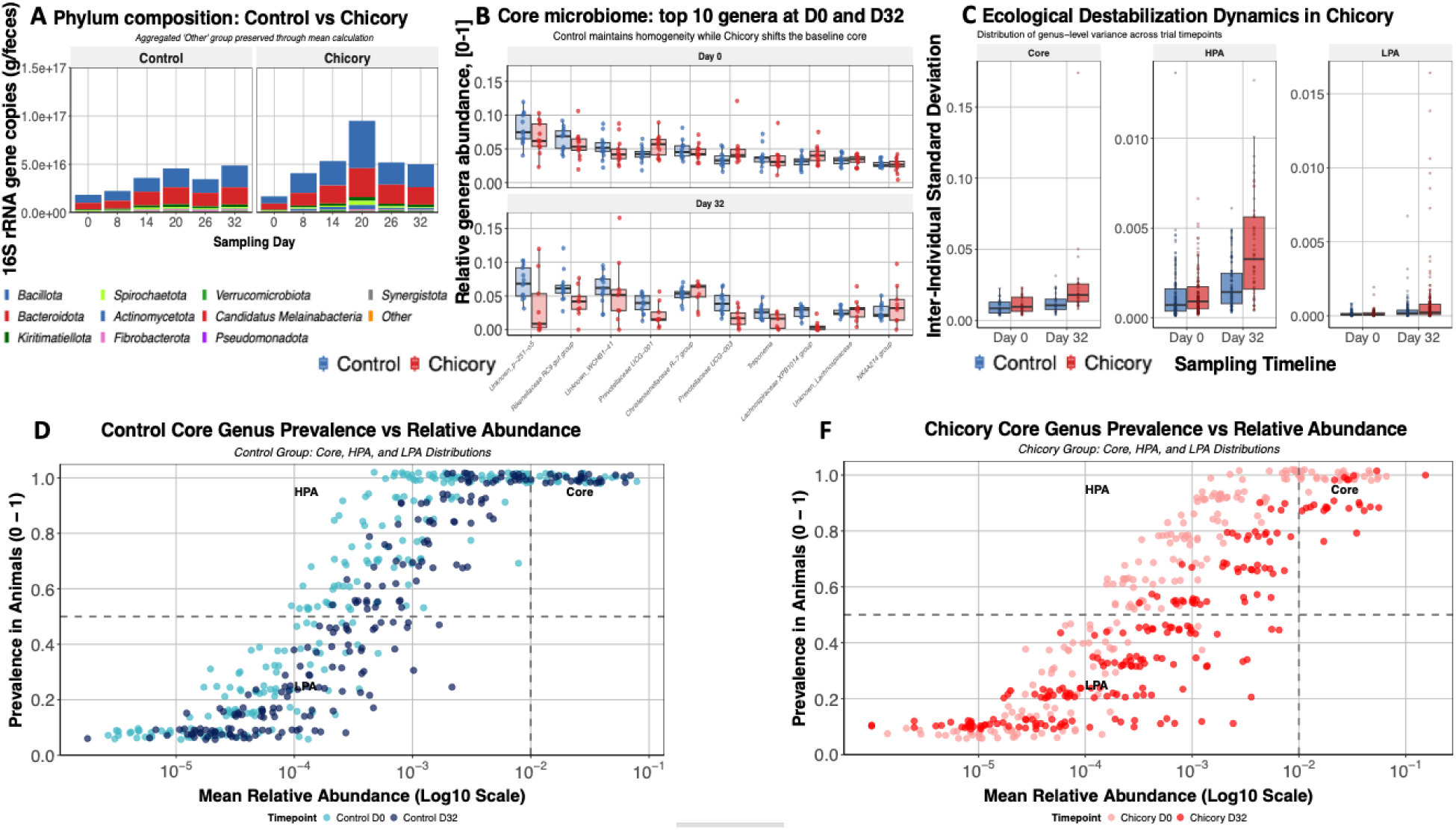
Microbial core community structure and ecological stability under chicory grazing. **(A)** Absolute phylum-level composition expressed as 16S□rRNA gene copies per gram of faeces across sampling days. Bacillot and Bacteroidota dominate both groups, while chicory grazing increases total bacterial load and amplifies fluctuations in minor phyla; **(B)** Core-microbiome composition showing the ten most abundant genera at Day□0 and Day□32. Control horses maintain a homogeneous baseline, whereas chicory grazing shifts relative abundances and broadens inter-individual variability; **(C)** Ecological destabilisation dynamics represented by genus-level inter-individual standard deviation for the core, the High-Prevalence Abundant (HPA), and the low-Prevalence Abundant (LPA) fractions. Variance remains low in controls but rises sharply in chicory-fed horses at Day□32, indicating reduced community stability; **(D–F)** Prevalence–abundance distributions for core genera in control (D) and chicory (F) groups. Each point represents a genus plotted by mean relative abundance (log□□ scale) and prevalence across animals. Core genera occupy the upper right quadrant (high prevalence an abundance), with HPA genera at intermediate levels and LPA genera at low prevalence and abundance. In controls, Day□0 and Day□32 points largely overlap, indicating a stable core structure. In contrast, chicory shows little overlap between timepoints, with several Day□0 core genera shifting toward lower prevalence or lower abundance at Day□32, expanding the LPA fraction and increasing dispersion within the HPA tier.

Beyond this broad taxonomic overview, the core microbiome architecture at Day 0 was highly conserved across horses (Figure 5B). The top ten core genera, including *Fibrobacter, Treponema,* and members of the *Prevotellaceae, Lachnospiraceae*, and *Rikenellaceae* families, were detected in nearly all individuals and exhibited low inter-individual variation, forming a stable High-Prevalence Abundant (HPA) backbone (Additional Table 6). Low-Prevalence Abundant (LPA) genera and rare taxa contributed minimally, indicating a coherent and homogeneous community structure at baseline (Figure 5C).

However, by Day □32, clear core differences emerged between groups. In the control horses, th core, HPA, and LPA fractions remained remarkably stable, with minimal shifts in prevalence or relative abundance and consistently low genus-level variance (Figure 5B). *Fibrobacter* and members of the *Lachnospiraceae* family (NK4A136 and XPB1014 groups) showed the highest persistence and lowest temporal fluctuation, together forming the backbone of the stable core in the control group. Inter-individual variance in the core fraction increased markedly between Chicory□D0 and Chicory□D32 (*P*□<□0.001), and between Control□D0 and Chicory□D32 (*P*□<□0.001), with still a significant difference between Control□D32 and Chicory□D32 (*P*□=□0.0018). The coefficient of core variation increased progressively in chicory-fed horses, while remaining relatively stable in controls, revealing a gradual loss of synchrony within the core microbiome. Statistical comparisons confirmed significant differences at Days□14,□26,□and□32 (*P*□=□0.0091, □0.0091, □and□0.0006). By the end of the trial, chicory-fed horses exhibited more than twice the median dispersion of controls (110□% vs□49□%), indicating that chicory grazing amplified inter-individual variability among core taxa. The HPA fraction exhibited parallel patterns, with all pairwise contrasts involving Chicory□D32 reaching high significance (*P* <□0.001; Figure 5C). Similarly, the LPA fraction showed strong divergence over time (*P*□<□0.001 for all comparisons), indicating that chicory grazing amplified inter-individual microbiota variability even among low-prevalence taxa (Figure 5D-F). Together, these results demonstrate that chicory not only modifies specific taxa but also reshapes the ecological scaffolding of the gut microbiome, transitioning it from a stable, shared core toward a more dynamic and individualised community structure.

#### Microbiota diversity and ecological resilience

To assess changes in gut microbial ecosystem complexity during the trial, we compared alpha diversity between groups using the Shannon index across all sampling days. At baseline, diversity did not differ between groups adj *P* > 0.05, Figure 6A). From Day□20 onward, however, the chicory group showed a significant decline in Shannon diversity (adj *P* = 0.029), with the effect strengthening by Day□32 (adj *P* = 0.001, Figure 6A). Consistent with this ecological signal, beta-diversity analyses further demonstrated a progressive divergence in community structure between groups over time. The PCoA plots based on Bray–Curtis’s dissimilarities showed substantial overlap at baseline, confirming that horses started the trial with comparable microbial communities. However, from Day□8 onward, samples from the chicory group began to separate gradually along the primary ordination axis, and this separation became more pronounced by Days□20–32. This PCoA pattern was supported by PERMANOVA results, which revealed no significant difference at Day□0 (R²□=□0.029, □*P*□=□0.580), but strong and consistent divergence thereafter: Day□8 (R²□=□0.258,□*P*□=□0.002), Day□14 (R²□=□0.278,□*P*□=□0.003), Day□20 (R²□=□0.408,□*P*□=□0.002), Day□26 (R²□=□0.299,□*P*□=□0.002), and Day□32 (R²□=□0.143,□*P*□=□0.005).

**Figure 6.**
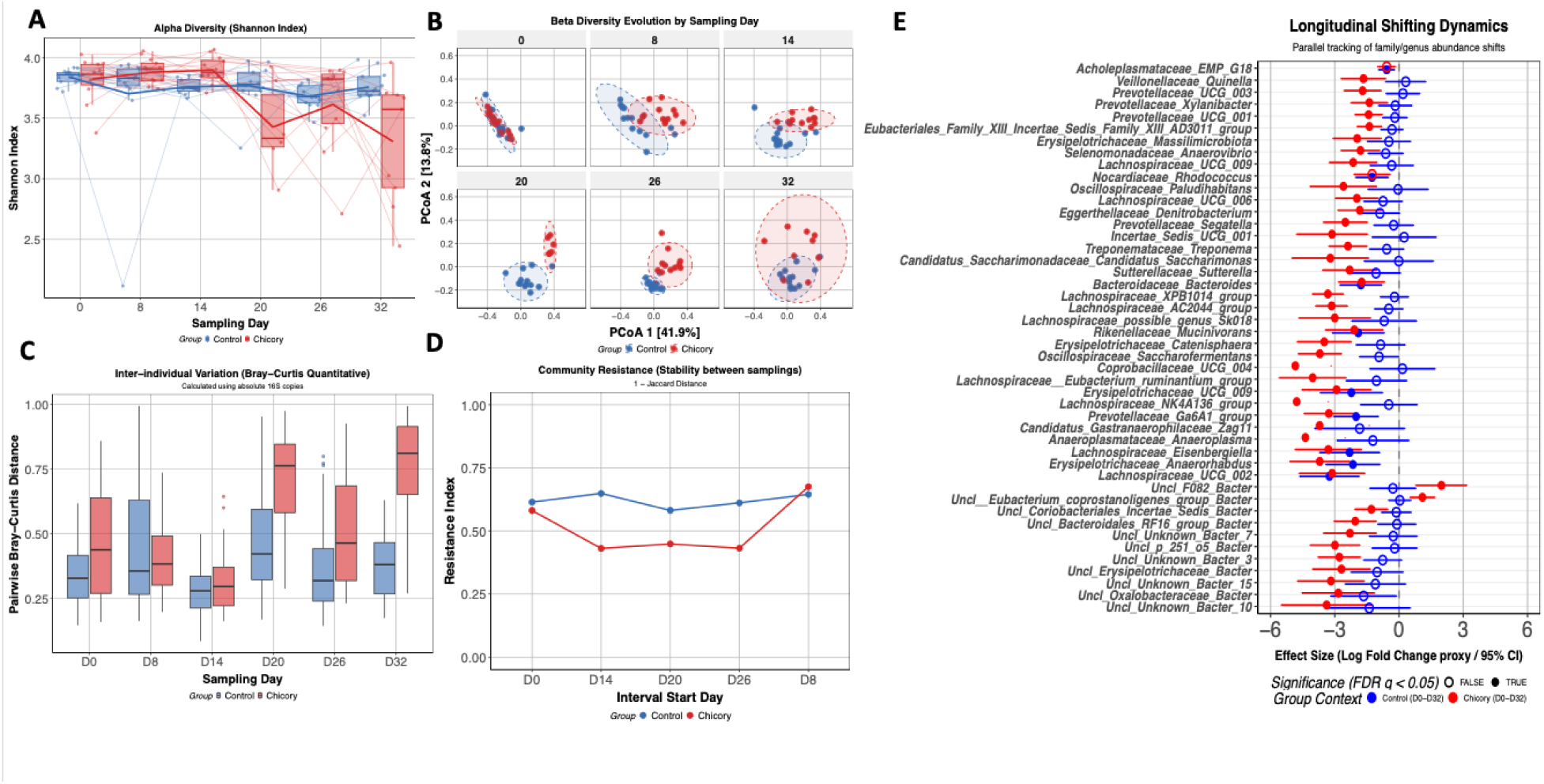
Microbial diversity, stability, and metabolic activity under chicory grazing. **(A)** Alpha diversity (Shannon index) showing temporal trends in microbial richness and evenness. Control horses maintained stable diversity, whereas chicory grazing induced greater variability and transient increases at mid-trial; **(B)**□Beta diversity (PCoA based on Bray–Curtis dissimilarities) illustrating progressive separation of chicor samples from controls, with clear divergence from Day□14 onward; **(C)**□Inter-individual variation (pairwise Bray–Curtis distances) calculated using absolute 16S□copies. Chicory grazing increased between-animal dissimilarity, indicating reduced community homogeneity; **(D)**□Community resistance (1□–□Jaccard distance) reflecting compositional stability between consecutive samplings. Chicory-fed horses exhibited lower resistance values, consistent with diminished temporal persistence; **(E)** This panel illustrates the genus abundance shifts between Day□0 and Day□32 for both experimental groups. Each row corresponds to a taxon, with horizontal bars representing the estimated effect size (log-fold-change proxy) and its 95□% confidence interval. Blue points indicate taxa whose abundance changed in the control group, while red points correspond to changes under chicor grazing. Filled circles mark statistically significant differences (adj *P*□<□0.05).

Analyses of interindividual variation using quantitative Bray–Curtis dissimilarities further confirmed the progressive divergence of microbial communities under chicory grazing. At baseline (Day□0), pairwise distances between individuals were already slightly lower in the chicory group than in controls (difference□=□–0.130□±□0.029,□*P*□<□0.001), indicating comparable initial homogeneity (Figure 6C). By Day□20, the chicory group exhibited markedly higher inter-individual dissimilarity, with significant differences at Day□20 (–0.224□±□0.043,□*P*□<□0.001), Day□26 (–0.156□±□0.031, □*P* <□0.001), and Day□32 (–0.377□±□0.035, □*P*□<□0.001; Figure 6C), whereas control horses maintained a compact community structure over time. Consistently, community resistance analysis revealed that chicory grazing reduced temporal microbial resistance. At baseline (D0) and early in the trial (D8), no differences were observed between groups (adj *P* = 0.375 and 0.898), indicating similar initial resilience. From D14 onward, resistance values declined markedly in the chicory group, with significant differences at D14 (estimate□=□0.282□±□0.070,□*P*□=□0.001), D20 (0.180□±□0.070,□*P* =□0.020), and D26 (0.192□±□0.060, *P* =□0.005). This reduction in compositional persistence suggests that chicory feeding weakened the community’s ability to maintain its structure over time, complementing the divergence and dispersion analyses that point to a transition towards a more unstable and individualised ecological state.

Beyond the global patterns of diversity and stability, longitudinal modelling revealed distinct taxonomic trajectories between the control and chicory groups. In control horses, only minor fluctuations were observed, mainly within *Lachnospiraceae* and *Prevotellaceae*, consistent with a stable fermentative core (Additional Table 7). In contrast, chicory grazing induced broad restructuring across several functional clusters (Additional Table 7). Within fibre-degrading and SCFA-producing families, genera such as *Anaerovibrio* (*Selenomonadaceae*), *Quinella* (*Veillonellaceae*), and *Xylanibacter* (*Prevotellaceae*) decreased markedly from D0 to D32. Members of *Erysipelotrichaceae, Oscillospiraceae*, and *Lachnospiraceae* (*UCG-002*, *UCG-006*, *UCG-009*, *Eisenbergiella*) also declined, suggesting a shift of rapid-turnover fermenters and mucin-associated taxa. Several uncultured lineages (e.g., *Candidatus Saccharimonas*, *Candidatus Gastranaerophilaceae Zag11*, and *Unknown Bacteria* clusters) also shifted significantly, indicating broader ecological remodelling beyond known genera (Figure 6E).

#### Microbial metabolic activity and environmental drivers of the microbial community structure

To better understand how chicory grazing reshaped the hindgut ecosystem, we combined microbial metabolic outputs with host- and environment-related variables within a unified ecological framework. All variables that significantly affected the host or gut environment were included in the *envfit* model, enabling us to identify the main drivers of shifts in microbial community structure. Short-chain fatty acids (SCFA) emerged as key contributors to community divergence. Acetate, butyrate, *iso*butyrate, valerate, and *iso*valerate showed strong, time-dependent correlations with the ordination axes, particularly during D14–D20, when chicory induced the most pronounced restructuring of the microbiota (Figure 7A-B). Thi pattern mirrors the transient increase in total SCFA concentrations observed in the chicory group. Total SCFA concentrations increased significantly in the chicory group at mid-trial (D14–D20; adj□*P*□= 0.002 and 0.037, respectively), reaching nearly twice the levels observed in controls before declining toward baseline by D32 (Figure□7C–D). Despite quantitative fluctuations, the relative SCFA composition remained dominated by acetate and propionate in both groups, with only a temporary enrichment in butyrate and *iso*butyrate at D14–D20 in the chicory group (Figure 7D; Additional Table 1).

**Figure 7.**
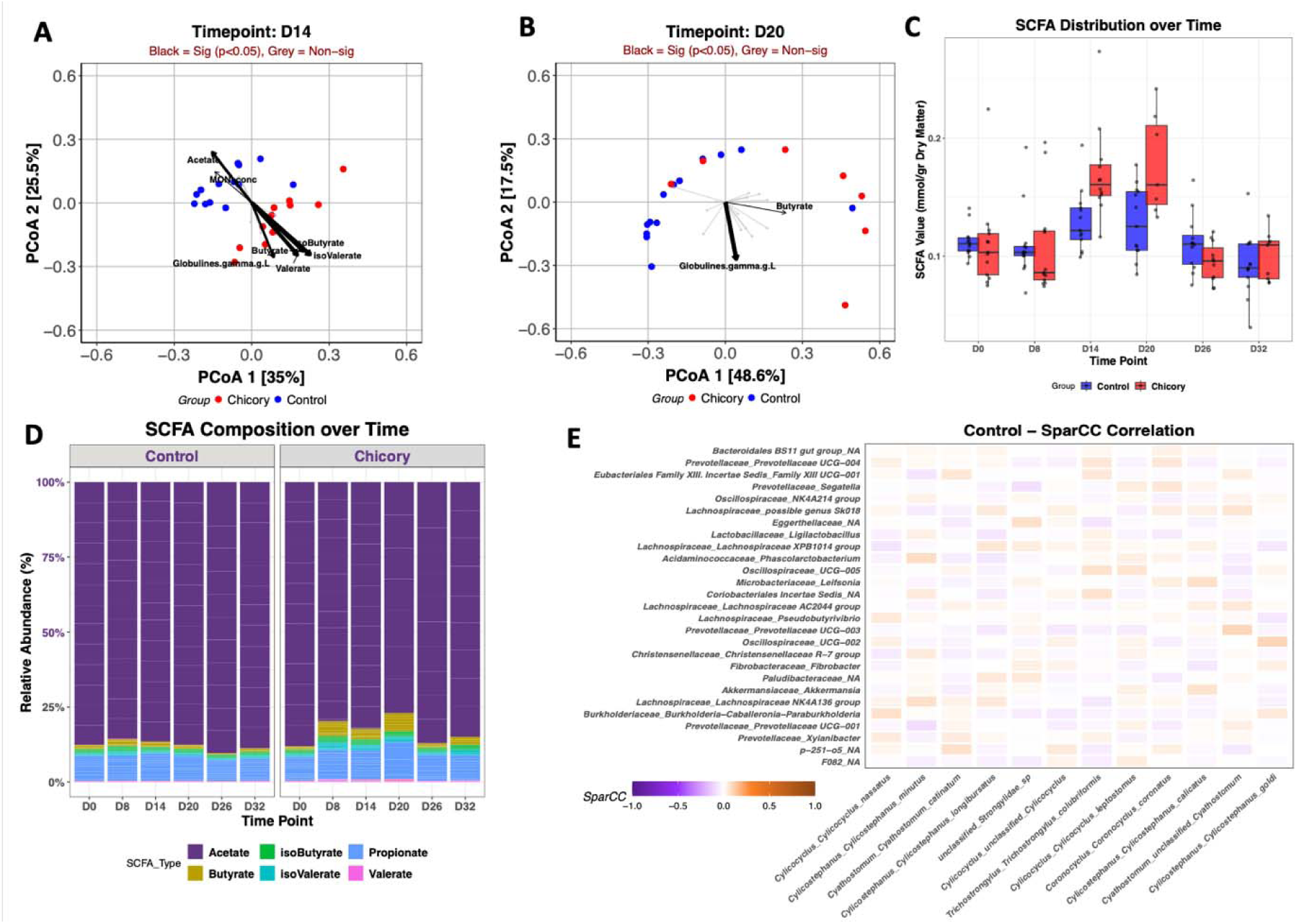
Short-chain fatty acid (SCFA) profiles and *envfit* associations with treatment over time. **(A-B)** Principal Coordinates Analysis (PCoA) of SCFA profiles at Days□14 and□20, with *envfit* vectors showing metabolites and serum variables significantly correlated with ordination axes (black, *p*□<□0.05; grey, non-significant). Chicory-fed horses (red) and controls (blue) **(C)**□Short-chain fatty acid (SCFA) concentrations over time (mmol/g□DM). Chicory grazing elevated total SCFA levels at Day□14–20, followed by a declin toward baseline; **(D)**□SCFA compositional profiles showing relative contributions of acetate, propionate, butyrate, and minor branched-chain acids. Despite transient quantitative increases, overall SCFA composition remained dominated by acetate and propionate in both groups. **(E)** Cross-domain correlation matrix (SparCC) for the control group, showing pairwise associations between dominant bacterial genera and nemabiome species. Rows represent bacterial taxa detected in faecal samples, and columns correspond to the main cyathostomin species identified in the nemabiome. Colour intensity indicates the SparCC correlation coefficient, ranging from negative (purple) to positive (red). Most correlations were weak (|r|□<□0.1, adjlJ*P*□>□0.05), reflecting limited direct co-variation between bacterial and parasitic communities.

In parallel, several host physiological markers, notably α2- and γ-globulins, neutrophil counts, and globulin concentration, also aligned significantly with microbial ordination patterns. These variables were especially influential at later time points (D20–D32), indicating that immune modulation and systemic responses covaried with microbial restructuring (Additional Figure 1).

### Weak microbiota–parasite co–variation despite strong chicory-driven nemabiome shift

To explore potential ecological interactions between the gut microbiota and the nemabiome, cross-domain correlations were computed using SparCC within each treatment group (Additional Table 7, Figure 7E). In both the control and chicory groups, the overall correlation structure was weak, with most coefficients close to zero (adj *P* > 0.05), indicating limited direct co-variation between bacterial and parasitic communities. Subtle patterns emerged that could help illustrate the functional triad linking host, microbiota, and parasite. In the control group, where the nemabiome remained largely stable and only *Cylicostephanus longibursatus* increased gradually over time, the correlation structure showed weak but coherent patterns: several commensal fermenters within *Lachnospiraceae* and *Prevotellaceae* displayed slight negative associations with the most persistent cyathostomin species, suggesting that these taxa may be less abundant under gut conditions that favour parasite persistence. Additionally, genera with known probiotic or antiparasitic potential, such as *Lactobacillus*, tended to show negative correlations with multiple nemabiome species. In contrast, under chicory grazing, where the nemabiome underwent a marked restructuring with strong declines in *Cylicocyclus ashworthi, C. leptostomus,* and *C. nassatus*, no clear or consistent correlation network emerged (Additional Table 8).

### Co-evolution of the Nemabiome and Microbiome

To explore potential co-evolutionary dynamics between the gut microbial and parasitic communities, we assessed the correspondence between microbial abundance profiles and nemabiome relative composition across all sampling days using Mantel and Procrustes analyses. Mantel tests revealed weak, non-significant correlations throughout the trial (r = –0.019 to 0.099, *P* > 0.10), indicating limited direct abundance-based similarity between the two communities. However, Procrustes analyses provided a more nuanced view of structural alignment. While early timepoints (D0–D20) showed high but non-significant concordance (M²□=□0.97–0.89,□*P*□>□0.10), significant associations emerged at D26 (M²□=□0.841, *P* =□0.045) and D32 (M²□=□0.827, *P* **=**□0.045), suggesting that the microbial and nemabiome ordinations became increasingly synchronised towards the end of the trial, likely indicative of a co-adaptive ecological response in which parasite community turnover and microbial reorganisation jointly shape the hindgut environment. To refine this interpretation, we also matched taxa at the genus level between the nemabiome and microbiome datasets. Genera showing consistent directional changes in both communities (e.g., *Cylicocyclus*, *Clostridium*, and *Lachnospiraceae* taxa) further support the hypothesis of functional co-evolution between microbial and parasitic assemblages under chicory grazing. Together, these findings suggest that the nemabiome and microbiome are not independent entities but dynamically interconnected systems, whose structural trajectories converge under dietary and ecological pressures.

## DISCUSSION

This study provides integrated evidence that grazing pastures dominated by chicory (80% of consumed bites on average) exert strong antiparasitic effects, reshape cyathostomin community structure, and induce a substantial reorganisation of the equine gut microbiota, while maintaining host behaviour, metabolic and immune homeostasis over the study period. These findings highlight the capacity of a single dietary intervention to simultaneously influence parasite dynamics and microbial ecosystem structure without overt compromise of host physiological stability.

A key outcome of this work was the pronounced reduction in faecal egg counts observed in horses grazing chicory-rich pastures. This result is consistent with our earlier work showing that chicory (*cv. Puna II*) can reduce cyathostomin egg excretion by 70–80% and impair larval development in grazing horses [50]. However, in contrast to our previous observations reporting impaired larval development [50], the present study revealed a higher larval development rate under chicory at specific time points, highlighting a dissociation between egg output and subsequent developmental success. This apparent discrepancy may be explained by the marked changes in the intestinal physicochemical environment induced by chicory. In the present study, chicory consumption was associated with a significant decrease in faecal redox potential, a transient acidification, and a shift in SCFA profiles, collectively indicating a transition toward a more reducing and fermentative hindgut environment. Comparable fermentation-driven changes have been observed in pigs, where dietary chicory increased butyrate-producing bacteria, altered SCFA concentrations, and lowered luminal pH [64]. Such fermentation-driven ecological restructuring is likely to have direct consequences for parasite biology. On the one hand, reduced redox potential and shifts in microbial fermentation may impair adult worm metabolism and fitness, thereby contributing to the observed reduction in FEC. On the other hand, these same environmental conditions may create a more favourable milieu for egg embryonation or larval development once eggs are excreted. For instance, reduced egg density under chicory may alleviate density-dependent constraints on development, while a more reducing and microbially active environment could enhance hatching success or larval viability. These indirect environmental effects may interact with chicory-derived bioactive compounds, particularly sesquiterpene lactones, which exhibit well-documented antiparasitic activity against adult stages and egg production in both equids and ruminants [49, 50, 54, 56, 65].

Importantly, despite these alterations in gut ecology, horses remained clinically stable, as evidenced by consistent body weight, normal metabolic indicators, and balanced immune profiles. These findings suggest that the antiparasitic effects of chicory do not induce measurable physiological stress in the host. They further support the conclusion that chicory acts directly on the parasites and/or modifies the immediate gut environment, rather than enhancing the host’s protective immune response. Behavioural observations also indicated comparable welfare between treated and control groups, with no changes in feeding behaviour or social interactions. A moderate increase in locomotor activity and feed intake was observed, which may plausibly reflect a reduced parasite burden and improved overall comfort.

At the nemabiome community level, the absence of a decline in overall diversity despite marked compositional shifts indicated that chicory did not act as a broad-spectrum suppressor, but rather imposed a selective ecological pressure on specific cyathostomin taxa. Relative abundance of dominant species such as *Cylicocyclus nassatus* [5, 14] declined to near background abundance under chicory, whereas the relative abundance of *Cylicostephanus calicatus* remained unchanged. Although *C. calicatus* is a common and frequently detected species in equine infections, current evidence does not support greater virulence than that of dominant taxa such as *C. nassatus* [5]. Its persistence under chicory exposure is more plausibly attributable to differential tolerance to dietary selective pressures, pointing to a reassembly of community structure rather than a shift in intrinsic pathogenicity.

Echoing the nemabiome reorganisation, the gut microbiota underwent an even more pronounced ecological transition. At baseline, microbial communities were highly homogeneous across individuals and dominated by typical hindgut fermenters, including members of the *Fibrobacter, Treponema*, and *Lachnospiraceae* lineages, which are consistently reported as core taxa in equine microbiomes [13, 14, 66–73]. Chicory grazing progressively disrupted this synchrony, leading to reduced Shannon diversity, increased inter-individual Bray–Curtis dissimilarity, and decreased temporal resistance. By Day□32, chicory-fed horses exhibited markedly higher dispersion than controls and a pronounced breakdown of the core microbiome, indicative of a transition from a stable, coordinated community to a more individualised and dynamic ecosystem.

This gut microbial shift likely reflects the combined influence of several interacting factors. First, chicory provides a distinct nutritional and biochemical profile, including low levels of DM and NDF but elevated levels of ADL, minerals, and bioactive compounds such as sesquiterpene lactones [74]. These components can alter both substrate availability and the physicochemical environment of the hindgut, thereby reshaping microbial niches [64]. Diet is a major determinant of equine microbiota structure, and even relatively subtle changes in fibre composition or lignification can induce individualised microbial responses [75, 76]. Similar patterns are observed in ruminants, where tannin- and sesquiterpene-rich forages such as chicory modulate rumen microbiota composition and fermentation processes, including shifts in functional pathways involved in fibre degradation and changes in SCFA profiles [77, 78]. Second, the concurrent reduction and restructuring of cyathostomin populations modifies host–parasite–microbiota interactions [14, 16, 26, 29], potentially altering immune signalling, mucosal function, and resource competition within the gut ecosystem. Helminths and microbes share the same intestinal niche [21] and interact both directly and indirectly through host immune pathways, forming a tripartite host–microbiota–parasite system. In horses, cyathostomin infections have been associated with shifts in microbial composition and community dynamics, although the direction and magnitude of diversity changes are inconsistent across studies [13–15, 22, 47, 79, 80]. Some reports describe modest reductions or no significant changes in alpha diversity, whereas others emphasise alterations in community structure or function rather than taxonomic diversity *per se*, indicating strong context dependency. In the present study, immune parameters remained largely stable despite nemabiome restructuring, with the exception of a change in IL-10, suggesting that systemic immune modulation was limited and likely pathway-specific. This constrained immune response further supports the interpretation that the pronounced decline in microbial diversity and resistance observed here reflects the combined effects of high chicory inclusion and substantial nemabiome remodelling, rather than a generalised immune-mediated disruption or a uniform response to either factor alone.

Whether such a marked loss of microbial diversity is detrimental remains an open question. From an ecological perspective, reduced diversity and lower resistance are often interpreted as indicators of decreased ecosystem robustness and an increased risk of dysbiosis, as diversity, redundancy, and network complexity are thought to underpin microbiome stability and resiliency [81, 82]. However, in the present study, the absence of clinical abnormalities, together with the maintenance of metabolic and immune homeostasis and the lack of negative behavioural indicators, suggests that, at least over the 32-day period, hosts were able to buffer this ecological reorganisation without apparent cost. This apparent tolerance is supported by emerging experimental evidence demonstrating that dietary fibre can directly modulate host physiology independently of microbiota composition. Recent work has shown that high levels of fermentable fibres, such as inulin, can alter host immune responses and infection susceptibility through pathways involving IL-18, IL-27, and tryptophan metabolism, while microbiota changes arise largely as downstream consequences of host-driven processes [83]. Taken together, these findings suggest that microbiome functionality, ecological redundancy, and host resilience likely play a more decisive role than microbial diversity and resistance alone in determining health outcomes. The transition toward a more individualised and less synchronised community observed in chicory-fed horses may therefore represent an alternative ecological state that remains compatible with host homeostasis over the short term.

## Conclusions

Chicory grazing at 80% inclusion in the bites clearly emerges as an effective ecological intervention that may reduce cyathostomin fitness and reshape the nemabiome in a species□specific manner, without compromising host metabolic or immune stability. These results reinforce chicory’s potential as a sustainable antiparasitic strategy and show that such a high level of inclusion can markedly alter microbial diversity and ecological resistance in horses. Importantly, the long-term consequences of this microbial destabilisation remain uncertain: reduced microbial diversity and resistance are not inherently detrimental, and it is unclear whether these shifts represent a transient adaptive response or the early stages of a new ecological configuration. Overall, chicory at 80% serves as both a natural anthelmintic and a strong modulator of gut ecology, highlighting the need for longer term functional studies to assess the health implications of these changes in the gut microbiota and microenvironment over extended grazing periods.

### Limitations

Although this study provides a high-resolution view of how chicory grazing reshapes parasite egg excretion, nemabiome structure, and gut microbial ecology, several limitations should be acknowledged to contextualise the findings. First, the study focused exclusively on cyathostomin strongyles, the dominant parasite group in grazing horses, but other gastrointestinal parasites—including *Strongyloides* spp.*, Oxyuris equi*, *Parascaris* spp., and tapeworms—may have been present at low levels and could have contributed to the observed ecological dynamics, particularly immune modulation or microbiota interactions. Because the nemabiome metabarcoding pipeline was optimised for *Cyathostominae* ITS-2 detection, leaving the full parasitic community and its potential influence on gut ecology only partially characterised. Second, the grazing period was intentionally restricted to 32 days to avoid reinfection within the cyathostomin prepatent period, but this short timeframe limits our ability to assess whether the observed microbial destabilisation and nemabiome restructuring persist, intensify, or resolve over longer periods. Third, the study involved naturally infected young horses with heterogeneous baseline nemabiomes, which reflect real-world conditions but introduce inter-individual variability that may mask subtler treatment effects. Fourth, although the chicory and control pastures differed primarily in botanical composition, other unmeasured environmental factors, such as microspatial grazing patterns, soil microbial communities, or fluctuations in plant secondary metabolites, may have contributed to the observed ecological shifts. Fifth, while the study documents a clear reduction in microbial diversity and ecological resistance under chicory, it does not establish whether these changes are functionally detrimental, beneficial, or neutral for long-term gut health, as functional metagenomics or metabolomics were not performed. Finally, the study focuses exclusively on young horses; whether adult horses, ponies, donkeys, or other equids would exhibit similar responses remains unknown, and cross-species comparisons with ruminants, where chicory often reduces parasite burdens but does not consistently reduce microbial diversity, suggest that host-specific digestive physiology may modulate the magnitude and direction of chicory’s ecological effects.

## MATERIAL AND METHODS

### Study Site and Ethical Approval

The study was conducted in September 2024 at the French Horse and Riding Institute (IFCE) Technical Platform in Chamberet, France (01°43′14″E, 45°35′03″N; 440 m a.s.l.). All procedures complied with the European Union Directive 2010/63/EU on the protection of animals used for scientific purposes. The experimental protocol received approval from the *Comité Régional d’Éthique pour l’Expérimentation Animale du Limousin* (CREEAL – CE033) and authorization from the French Ministry of Research (protocol APAFIS#51175-2024100710358148).

### Pasture Characteristics and Agronomic Management

The Chamberet site is characterised by an oceanic climate and an episkeletic podzol soil type, as described in the European Soil Database (Joint Research Centre, EU; http://eusoils.jrc.ec.europa.eu/). Two pasture types were used during the trial: a chicory-based sward (*Cichorium intybus*, Puna II variety) and a permanent grassland serving as the control (Figure 1BC). The permanent pasture was composed primarily of *Festuca arundinacea*, *Phleum pratense*, *Poa abbreviata*, *Holcus lanatus*, and *Dactylis glomerata*. The chicory pasture consisted of two plots of comparable size: one (named 13I) was sown on 22 April 2024 at a rate of 9 kg/ha using the Puna II variety; the second (16I), originally established on 11 April 2022, was oversown on 28 May 2024 at the same seeding rate. The permanent grassland (13II) covered 3.9□ha, whereas the chicory treatment relied on 4.1 ha of sown pasture. Each pasture was divided into five sub-plots (named ‘a’ to ‘e’; approximately 0.80 ha each), and horses grazed them in rotation on a fixed schedule of three days per sub-plot. The Control group grazed subplots 13IIa, 13IIb, 13IIc, 13IId, and 13IIe successively before starting again, while the Chicory group grazed the subplots in the following order: 13Ia, 13Ib, 16Ia,16Ib and 13Ic. From 26 September (e.g. for the last five days of the study), horses in both groups were offered three sub-plots at the same time as the amount of available food resources became limited. This rotational design ensured comparable grazing pressure across treatments and minimised confounding effects related to pasture heterogeneity [84]. Water was available *ad libitum* in both pastures throughout the study.

Fertilisation practices differed markedly among the three experimental plots (chicory pasture: 13I and 16I, permanent grassland: 13II), reflecting their distinct agronomic histories and expected botanical dynamics. The chicory-based plot 13I received 2□t/ha of horse manure on 12 September 2023, followed by 100□kg/ha of ammonium nitrate on 13 May 2024 to support chicory establishment and maintain adequate forage biomass during grazing. Plot 16I, which also contained chicory, received 2□t/ha of manure earlier in the year, on 6 January 2023, but no mineral fertilisation in 2024. In contrast, the permanent pasture (13II; 3.9□ha) had not received any organic or mineral amendments in recent years, consistent with its long-term management as an unfertilized grassland. These contrasting fertilisation regimes were carefully documented because they influence soil nutrient availability, plant growth rates, secondary metabolite production, and ultimately the grazing behaviour and dietary choices of the young horses. They also provide essential context for interpreting potential differences in forage structure, chemical composition, and exposure to bioactive compounds such as sesquiterpene lactones.

### Animals and Pre-Trial Management

Twenty-six naturally infected, between 1- and 2-year-old Anglo-Arabian saddle young horses (e.g., 20 two-year-olds and 6 one-year-olds) were enrolled in the study. All animals had remained undrenched for 137 days prior to the trial, with their last anthelmintic treatment administered on 24 April 2024 (200 μg ivermectin/kg body weight and 1 mg praziquantel/kg body weight; EQVALAN® DUO, France).

Horses were allocated to treatment groups balanced for sex (seven to eight females and five to six geldings per group), age (10 two-year-old and 3 one-year-old horses in each treatment), faecal egg counts (FEC) measured on 19 August (chicory group 1020 ± 842 eggs per gram of wet faeces; control group 1067 ± 837 eggs per gram of wet faeces), and body weight measured on 26 August (421.2 ± 33.1 kg versus 418.8 ± 29.7 kg). This ensured that initial parasitological and physiological conditions were comparable between groups.

None of the horses received antibiotic therapy during the sampling period, and diarrhoea was not detected in any animal.

### Experimental Design

The grazing trial lasted 32 days, from 29 August to 30 September 2024, a duration intentionally chosen to remain shorter than the pre-patent period of cyathostomes and thus prevent reinfestation from newly deposited eggs. The 26 horses were divided into two balanced groups of 13 individuals. The overall experimental design is summarised in Figure 1A.

### Sampling and Parasitological Analyses

Faecal and blood samples were collected simultaneously from each of the 26 individuals at six different time points: day 0 (D0), day 8 (D8), day 14 (D14), day 20 (D20), day 26 (D26), and day 32 (D32). Blood samples were taken while the animals were restrained in a squeeze chute. The same investigator examined the horses at each sampling point for body weight, body temperature, and general state. Body weight was measured using an electronic scale (PUEC31, Radwad Electronics, Poland).

Faecal samples were collected non-invasively from the ground following direct observation of defecation. Because the animals were young, no rectal sampling was performed to avoid unnecessary handling stress. Immediately after deposition, faecal balls were inspected, and only those not in contact with the ground were selected to minimise environmental contamination. Single-use gloves and sterile sampling tubes were used to collect faecal samples from each animal at each time point.

For each sample, material intended for microbial and biochemical analyses was collected by aseptically sampling the inner portion of the faecal ball, thereby avoiding the external surface exposed to air or potential contaminants. This sampling strategy follows the procedures previously used by Mach et al., ensuring consistency with established methodologies for equine faecal microbiota and metabolite studies [66, 85, 86]. Faecal samples were subdivided immediately for parasitological, microbiological, and chemical analyses. Precisely, 1 g faecal sample was snap-frozen and stored at -80 °C for microbial DNA extraction.

FECs were determined using a modified Mini-FLOTAC technique. Briefly, 3 g of faecal material were homogenised in 27 mL of water. The suspension was filtered, transferred to a centrifuge tube, and centrifuged. After discarding the supernatant, the sediment was resuspended in a saturated sodium chloride flotation solution (specific gravity = 1.18–1.20). The homogenised suspension was then used to fill both chambers of a Mini-FLOTAC device [87]. Following a 10-minute flotation period, gastrointestinal strongyle eggs were enumerated in both chambers, yielding a theoretical detection limit of 5 eggs per gram of faeces.

Coprocultures were performed to allow the development of nematode eggs to the infective third-stage larvae (L3). Approximately 30–50 g of faeces per individual were mixed with vermiculite and incubated in large Petri dishes at 24 ± 1 °C for 12–15 days, with regular aeration and moisture maintenance. Infective L3 larvae were subsequently recovered using a Baermann apparatus with tap water. Recovered larvae were frozen at –80°C for subsequent DNA extraction.

At the lab, faecal pH and redox potentials were immediately determined after 10% faecal suspension (wt/vol) in saline solution (0.15 M NaCl solution) with a multi-parameter portable meter (Multi 3620 IDS, WTW, GmbH, Weilheim, Germany) equipped with field redox and pH electrodes (SenTix, WTW, Weilheim, Germany). Briefly, 1 g of faecal samples was diluted in 10 mL of distilled water and centrifuged at 8,000 rpm for 10 min.

Dry matter (DM) content was determined for each faecal sample. Briefly, approximately 2□g of fresh faeces were weighed, dried at 60□°C for 48□h to a constant mass, and reweighed. Dry matter percentage was calculated as: DM (%) = (dry weight / fresh weight) × 100. FEC and SCFA concentrations obtained from gas chromatography were subsequently corrected for DM content.

For SCFA extraction, 1□g of faecal material was mixed with 0.2□mL of sulfuric acid (25% v/v) and stored at −20□°C until analysis. Faecal SCFA concentrations were measured as previously described [88]. Faecal samples and sulfuric acid mixes were thawed on ice overnight for gas chromatography analysis. These samples were centrifuged at 2,880 × g for 20 min at 4 °C to separate the liquid phase. For protein removal, 1 mL of supernatant was mixed with 200 μL of 25% metaphosphoric acid (v/v) and centrifuged at 20,000 × g for 15 min at 4 °C. Then, 100 μL of the supernatant was added to 75 μL of 4-methylvaleric acid (0.2% v/v), an internal standard, and 900 μL of ultrapure water. From this mixture, 1□μL was then injected into a gas chromatograph (Hewlett Packard, Model 7890A) equipped with a DB-FFAP column (30□m□×□0.53□mm i.d., 1-μm film thickness, Agilent Technologies, Palo Alto, CA, United States) and an FID detector (Avondale, PA, United States). Chromatograms were integrated using Chromeleon software (Thermo Fisher Scientific, version 6.8, Waltham, MA, United States). Six SCFAs, acetic acid (C2:0), propionic acid (C3:0), butyric acid (C4:0), valeric acid (C5:0), *iso*butyric acid (*iso*-C4:0), and *iso*valeric acid (*iso*-C5:0), were quantified using automated gas separation, according to the method of Playne (1985) [89]and modified as follows. The sum of the six SCFA concentrations was defined as the total concentration and was used to obtain the molar proportions of each SCFA. The proportion or relative abundance of each fatty acid was calculated by dividing the concentration of each fatty acid by the total fatty acid content.

At each sampling time point, photographs of freshly deposited faeces were systematically taken to document their macroscopic characteristics, allowing a standardised assessment of consistency and colour for each animal throughout the study (Additional Figure 2).

### Blood Sampling and Biochemical Analyses

Blood samples were collected at each sampling point into EDTA-K3 tubes for haematology, heparin tubes for biochemical analyses, and PAXgene tubes for immune gene expression profiling. Haematological parameters, including leukocyte subsets, erythrocytes, hematocrit, mean corpuscular volume, and thrombocytes, were quantified using an MS9-5 Haematology Counter® digital automatic haematology analyser (Melet Schloesing Laboratories, Osny, France).

Serum biochemical parameters, including albumin, globulin, total proteins, and urea, were measured using colourimetric assays on an M-Scan II analyser (Melet Schloesing Laboratoires, Osny, France). TNFα and IL-10 were measured in serum samples by ELISA using paired equine capture and detection antibodies (DuoSets, R&D Systems, UK) according to the manufacturer’s instructions. ELISA OD (450 nm) values were obtained using a Multiskan FC plate reader (Thermo Fisher, USA). Detection limits for TNFα and IL-10 were 31.2 pg/mL and 312 pg/mL, respectively. Samples below the minimum range were assigned an arbitrary value of half the detection limit. Adaptive immune responses were evaluated by quantifying TH2-associated mRNA expression in PAXgene-stabilised blood samples, including IL-13, IL-4, and IL-5 (Additional Table 9).

Total RNA was extracted from PAXgene whole blood samples using the PAXgene Blood RNA Kit (Cat. No. 762174, QIAGEN). RNA concentration and purity were assessed using a NanoDrop 2000c spectrophotometer (Thermo Scientific). Subsequently, 270 ng of total RNA was reverse-transcribed into cDNA using the SuperScript® VILO™ cDNA Synthesis Kit (Cat. No. 11754250, Life Technologies), according to the manufacturer’s instructions. Relative gene expression of IL-4, IL-5, and IL-13 was quantified by qRT-PCR on a QuantStudio™ 12K Flex Real-Time PCR System (Life Technologies), and all conditions were run in duplicate. Each reaction was processed in a total volume of 25 µl containing 2X Taqman® Universal PCR Master Mix (Cat. No. 4305719, Life Technologies), primers, probes, and 2.5 µL of cDNA diluted at 1:2 (Additional Table 9). The thermal cycling conditions were as follows: an initial denaturation step of 10 min at 95°C, followed by 50 cycles consisting of 15 s at 95°C and 1 min at 60°C. Negative controls and calibrators were included, and relative expression was normalised to the reference genes GAPDH and ACTB using the 2^-ΔΔ^Ct^method [90]. Analysis was performed on the Thermo Fisher Connect Platform. Gene expression results were expressed as log_2_ fold changes, using the mean expression values of all control horses at day 0 as the calibrator reference.

### Vegetation and dietary choices measurements

In each treatment, the distribution of vegetation heights (100 sample points per diagonal, measured at the first place where a stick contacted the sward surface) was assessed six times during the experiment, on the second or third day of the presence of horses in a subplot (D1, D7, D13, D19, D25, and D31). At each sample point, we also recorded available bites characterised by the dominant (e.g., > 50%) item among chicory, grasses, legumes, forbs or plant species with anthelmintic properties reported in small ruminants (e.g., *Plantago lanceolata, Rumex obtusifolius* L., and *Chenopodium album* L). Chicory and plants with anthelmintic properties were recorded as present if they were observed, but not dominant. When no vegetation was present, the categories “bare soil” or “faeces” were recorded.

On the same six days, dietary choices were measured by two observers (one in each group) using scan sampling method with activity recorded in both groups every 15 min for each animal from 08:00 to 12:00 and from 15:30 to 19:30. For each grazing horse, the observer moved as close to the animal as possible (∼2m) without disturbing it to record the botanical composition of one selected bite, distinguishing chicory, grasses, legumes, forbs, and anthelmintic species. Observers decided beforehand which bite to register (e.g., the fifth one) to avoid bias in picking the most visible bites. The observer also recorded the ear position during grazing (e.g., forward, neutral, back), as an ear-back state is a potential sign of discomfort. These observations required animals to be trained for a week before measurements.

To complement these behavioural observations, we also characterized the quality of available vegetation by analysing dry matter (60°C to constant weight), crude protein (Dumas method N×6.25) and fiber content (NDF, ADF and ADL [91]) in one composite sample per treatment (from six 10cm × 5m-long strips cut to ground level) at D1, D7, D12, D19, D25, D32. The same analyses were performed on a composite sample from the permanent grassland where the animals had grazed the day before the trial.

### Weather Data

We use daily weather station observations from *Météo* France, the French national meteorological service (https://meteofrance.fr). All the observations were made at a station 5 km from the experimental farm. The stations calculate daily T_min_ as the lowest T_a_ observed from 0 coordinated universal time (UTC) on the previous day until 0 UTC, and daily T_max_ as the highest T_a_ observed from 0 UTC on the day until 0 UTC the following day. The T_mean_ was calculated as the mean of all (at least 24) T_a_ observations from 0 UTC on the day until 0 UTC the next day. The weather and altitude conditions remained constant for all animals throughout the study. These data were expressed as daily averages and used to contextualise grazing conditions and potential environmental influences on parasite development and plant chemistry.

### Behavioural observations

Behavioural observations were conducted to characterise the daily activity patterns, social interactions, and overall welfare of the horses during the grazing trial. Direct observations were conducted every 15 minutes on each horse during two daily periods: from 08:00 to 12:00 in the morning and from 15:30 to 19:30 in the afternoon, for a total of 32 observations per individual over six observation days using an instantaneous scan sampling method (see [92] for a detailed description of the methodology). This resulted in a total of 192 scans per horse across the study. For each animal, we recorded a comprehensive set of behaviours, including foraging activity (e.g. biting, chewing or swallowing grass, or walking with its muzzle close to the sward), locomotion (e.g. at least three successive steps with the head raised; one steps corresponds to the movement of one front limb), standing, stillness (standing with a hindquarter resting on the hoof toe, or lying), and social interactions. Social behaviours were classified as positive, negative, or neutral, thereby capturing the quality and frequency of affiliative and agonistic interactions within each group.

We also systematically recorded abnormal or welfare-related behaviours during scans, such as indicators of anxiety, apathy, stereotypic behaviours, and aggression directed either toward conspecifics or toward humans. Definitions of behavioural categories, social interactions, and welfare-related behaviours were based on previously published ethograms and observational protocols described by the Lansade group [67, 93–95]. This observational framework provided an integrated overview of the animals’ behavioural repertoire and social dynamics, enabling us to assess how grazing conditions and dietary treatments may influence their behaviour (social and non-social) and well-being. Finally, the horses’ daily grazing time was recorded over 24 h using Ethosys® collars (Greenway System GmbH, Frankfurt, Germany) [96]. The horses were acclimated to the collars for 1 week before the experiment began.

### Statistical analysis for physiological and parasitological variables

Statistical analyses were performed using appropriate software (e.g., R). Physiological, biochemical, and parasitological parameters were analysed using a combination of Wilcoxon rank-sum tests, Welch’s t-tests, or linear mixed-effects models (LMER), depending on the distribution and structure of each variable. For instance, FEC, which did not meet the assumption of normality, was compared between the chicory and control groups at individual time points using the Wilcoxon rank-sum test. Larval development data were analysed using Welch’s t-test to account for unequal variances between groups, with test statistics (*t*-values, degrees of freedom) and corresponding *p*-values reported. Physicochemical parameters, including faecal redox potential and pH, were described over time between treatments using appropriate models, including mixed-effects models for repeated measures, and statistical significance was assessed using *p*-values.

### Microbiota study and statistical analysis

#### Faecal microbial 16S rRNA gene sequencing data production and analysis

Faecal samples were thawed on ice to isolate faecal DNA. Total DNA was then extracted using the EZNA Stool DNA Kit (Omega Bio-Tek, Norcross, Georgia, USA) according to the manufacturer’s instructions.

Faecal DNA purity and concentration were estimated using a NanoDrop Spectrophotometer (NanoDrop Technologies Inc., Wilmington, DE, USA) and a Qubit 2.0 fluorimeter (dsDNA HS assay kit; Thermo Fisher). In all cases, negative control samples were included alongside biological samples during DNA extraction to minimise DNA contamination prior to sequencing. Additionally, contamination was minimised through laboratory techniques such as UV irradiation of the material, use of ultrapure water, use of DNA-free Taq DNA polymerase, and separation of pre- and post-PCR areas. We also included ZymoBIOMICS™ Microbial Community Standard bacterial cells (Catalogue No. D6300, Zymo Research Corp., Irvine, CA, United States) (“Zymobiomics-Cells”) as a positive control during DNA extraction. According to the manufacturer’s specifications, Zymobiomics-Cells consist of three gram-negative bacteria within the phylum Proteobacteria [*Pseudomonas aeruginosa* (theoretical composition in terms of 16S rRNA gene abundance: 4.2%), *Escherichia coli* (10.1%), and *Salmonella enterica* (10.4%)] and five gram-positive bacteria within the phylum Firmicutes [*Lactobacillus fermentum* (18.4%), *Enterococcus faecalis* (9.9%), *Staphylococcus aureus* (15.5%), *Listeria monocytogenes* (14.1%), and *Bacillus subtilis* (17.4%)]. All samples were diluted to 10 ng/µL and sent to the sequencing facility.

The V3-V4 hyper-variable region of the 16S rRNA gene was amplified, as previously reported by our team [13, 14, 66, 86]. The 16S rRNA gene libraries were prepared according to Illumina’s protocol (# 15044223 RevB, Illumina Inc., San Diego, CA, United States). To minimise the risk of cross-contamination and pipetting errors, the workflow was automated using a high-throughput liquid handler, the Freedom EVO NGS workstation (TECAN). Briefly, V3-V4 hypervariable regions were first amplified from 10 ng of genomic DNA using the following primers: (i) Forward Primer: TCGTCGGCAGCGTCAGATGTGTATAAGAGACAGCCTACGGGAGGCAGCAG and (ii) Reverse Primer: GTCTCGTGGGCTCGGAGATGTGTATAAGAGACAGGACTACHVGGGTATCTAATCC and 2X KAPA HiFi HotStart ReadyMix (Kapa Biosystems). PCR cycle conditions were 95 °C for 3 min, 25 cycles of (95 °C for 30 s, 55 °C for 30 s, 72 °C for 30 s), and then a final extension of 72 °C for 5 min. The libraries were purified using AMPure XP beads (Beckman Coulter). Dual indexes and sequencing adapters from the Illumina Nextera XT index kits (Illumina) were added in a second PCR using 2× KAPA HiFi HotStart ReadyMix (Kapa Biosystems). Cycle conditions were 95 °C for 3 min, 8 cycles of (95 °C for 30 s, 55 °C for 30 s, 72 °C for 30 s), then a final extension of 72 °C for 5 min. Ready-to-sequence libraries were purified using AMPure XP beads (Beckman Coulter) and quantified by fluorescence using the QuantiFluor One dsDNA kit (#E4870, Promega) on the GloMax system (Promega). Quality control was performed using a 2200 TapeStation system with the DNA 1000 ScreenTape (# 5067-5582, Agilent). The library pool was quantified by qPCR with the KAPA Library Quantification Kit for Illumina platforms (Kapa Biosystems). Sequencing was performed on a MiSeq system (Illumina) with the MiSeq Reagent v3 kit (600 cycles) in a 2□×□250 bp mode. The mock community DNA sample, “20 Strain Staggered Mix Genomic Material” (ATCC® MSA-1003TM), was used to control sequencing errors at concentrations ranging from 6.08 ng/µL to 10.14 ng/µL. The mock community declared components are: 0.18% *Acinetobacter baumannii* (ATCC 17978), 0.02% *Actinomyces odontolyticus* (ATCC 17982), 1.80% *Bacillus cereus* (ATCC 10987), 0.02% *Bacteroides vulgatus* (ATCC 8482), 0.02% *Bifidobacterium adolescentis* (ATCC 15703), 1.80% *Clostridium beijerinckii* (ATCC 35702), 0.18% *Cutibacterium acnes* (ATCC 11828), 0.02% *Deinococcus radiodurans* (ATCC BAA-816), 0.02% *Enterococcus faecalis* (ATCC 47077), 18.0% *Escherichia coli* (ATCC 700926), 0.18% *Helicobacter pylori* (ATCC 700392), 0.18% *Lactobacillus gasseri* (ATCC 33323), 0.18% *Neisseria meningitidis* (ATCC BAA-335), 18.0% *Porphyromonas gingivalis* (ATCC 33277), 1.80% *Pseudomonas aeruginosa* (ATCC 9027), 18.0% *Rhodobacter sphaeroides* (ATCC 17029), 1.80% *Staphylococcus aureus* (ATCC BAA-1556), 18.0% *Staphylococcus epidermidis (*ATCC 12228), 1.80% *Streptococcus agalactiae* (ATCC BAA-611), 18.0% *Streptococcus mutans* (ATCC 700610).

A subsequent amplification was necessary to index sequences on the sequencing cell (NexteraXT Index Kit, FC-131–1001/FC-131–1002).

#### Processing of 16S rRNA gene sequencing

The quality of the reads was verified using FASTQC (v. 0.12.1) and MultiQC (v. 1.27.1). Adapter trimming and quality filtering were performed with BBMap/BBDuk v39.18 using the parameters ktrim=r, k=23, mink=11, hdist=1, tbo, tpe, minlength=50, ziplevel=6, after which FastQC and MultiQC were rerun to confirm complete removal of adapters. Amplicon sequence variants (ASVs) were inferred using the DADA2 pipeline (R package dada2), applying filterAndTrim(maxN=0, truncQ=2, trimLeft=10, truncLen=c(250,250), maxEE=c(2,2), rm.phix=TRUE, compress=TRUE, multithread=TRUE) followed by error-model learning, denoising, merging of paired reads, and chimaera removal. Taxonomic assignment was performed using SILVA v138.2 (*silva_nr99_v138.2_toGenus_trainset.fa.gz* and *silva_v138.2_assignSpecies.fa.gz*). DADA2 initially identified 11,007 ASVs, each representing a unique denoised DNA sequence. ASVs lacking taxonomy at all ranks, eukaryotic contaminants (Mitochondria, Chloroplast), and non-target environmental phyla (Planctomycota, Cyanobacteriota, Thermodesulfobacteriota, Halobacteriota, Nitrospirota, Nitrospinota) were removed. Because multiple ASVs frequently represent the same biological taxon, owing to intraspecific 16S rRNA variation, the presence of several slightly divergent 16S operons within a single genome, and the tendency of reference databases to collapse closely related sequences when taxonomic resolution is limited. ASVs were subsequently collapsed into taxonomic lineages following the approach described by Bars-Cortina et al [97]. As a final harmonisation step, lineage assignments were updated and standardised according to the NCBI Taxonomy database (ftp.ncbi.nlm.nih.gov/pub/taxonomy/;10/08/2025) using the *taxonomizr* R package, ensuring consistent and up-to-date taxonomic nomenclature across the dataset.t

The phyloseq (v. 1.54.0) [98], vegan (v. 2.7.2) [99] and microbiome packages (v. 1.32.0; http://microbiome.github.io) were used in RStudio (v. 2023.09.03) for downstream analysis. Data were aggregated at the genus, family, order, class, and phylum levels using the taxonomic-agglomeration method in the phyloseq R package, which merges taxa at a user-specified taxonomic level. To assess and mitigate potential contamination in sequencing data, we employed the decontam (v. 1.22.0) R package to identify and visualise possible contaminant ASV features in negative control samples. The *isContaminant ()* function was applied using the frequency-based method, which models the prevalence of each ASV as a function of the input DNA concentration. ASVs statistically classified as contaminants (*P* < 0.05) were excluded from downstream analyses. To further reduce the likelihood of false-positive taxonomic assignments arising from misalignment or low-level background noise, we applied an abundance threshold of 0.01% using the *filterfun_sample()* function from the phyloseq R package, thereby retaining only biologically relevant taxa. To validate sequencing accuracy and compositional fidelity, we analysed mock community samples using the ChkMocks R package (v.0.1.4). The *checkZymobiomics()* function was used to compute Spearman’s rank correlation between observed and theoretical community profiles. We confirmed that negative controls consistently failed to produce visible bands on agarose gel electrophoresis, and purified amplicon concentrations were below the detection threshold (< 1 ng/μL), supporting the absence of amplifiable DNA contamination.

#### Microbiota diversity and richness analysis: **α**- and **β**-diversity, core and accessory microbiota

The microbiome R package generated measures of evenness, dominance, divergence, and abundance indexes.

The gut α-diversity indices between treatment and time points were compared using a generalised linear model (*glmer()* function of the lme4 R package [100] (v. 1.1.38)), with day and treatment included as fixed effects and individuals as a random effect. Comparisons of α-diversity parameters between treatments were performed using the Kruskal-Wallis test, a nonparametric test for comparing the medians of three or more independent groups, with the function *kruskal.test()* from the stats R package.

Bray-Curtis dissimilarity was calculated using the phyloseq R package to estimate β-diversity. β-diversity was visualised using non-metric multidimensional scaling (NMDS) in the vegan R package, specifically the *metaMDS()* function. Stress values were calculated to determine the dimensions for each NMDS.

To assess the influence of treatment and time on microbial community structure, we performed a permutational multivariate analysis of variance (PerMANOVA) using Bray–Curtis dissimilarity. ASV tables were extracted from the phyloseq objects and converted to distance matrices using the *distance()* function from the phyloseq package *(method = “bray”)*. PerMANOVA was conducted using the *adonis2()* function from the vegan package, with Treatment and Time_point_day as explanatory variables and Animal_ID as strata to account for repeated measures. The model was run with 10,000 permutations to assess statistical significance.

#### Microbial Core Ecological Dynamics Analysis: three-layered landscape

Genera were classified into ecological categories based on these thresholds for relative abundance and prevalence. Taxa with a mean relative abundance ≥ 1% and prevalence ≥ 50% were classified as Core taxa. Taxa with a mean relative abundance < 1% but prevalence ≥ 50% were classified as High-Prevalence Low-Abundance (HPA) taxa, representing widespread but low-abundance community members. Taxa with both mean relative abundance < 1% and prevalence <50% were classified as Low-Prevalence Low-Abundance (LPA) taxa, representing rare or infrequently detected genera. Taxa exhibiting high abundance, but low prevalence, were retained as a separate auxiliary category when present.

The ecological category was quantified for each treatment and time point. Distributions were visualised using prevalence – abundance scatter plots. Relative abundance and prevalence thresholds were used to visually determine the frontier between ecological categories. Time points were used to visualise the dynamics of the categories over time. For each of these three-layered landscapes, inter-individual variability distributions were compared between groups and time points using the standard deviation of genus-level relative abundances. The variance distribution was visualised by comparing observations of individual genera at each core, HPA, and LPA.

Pairwise comparison of inter-individual variances of the different groups was performed independently of each ecological category using the Wilcoxon rank-sum test. The resulting p-values were corrected for multiple testing using the FDR procedure and considered statistically significant.

Moreover, longitudinal dynamics of the core microbiome members were analysed to identify the most stable members within the microbial communities. For each treatment group and time point, the median relative abundance (MAD) and dispersion were calculated at the genus level. Dispersion was estimated using the median absolute deviation (MAD). Subsequently, the coefficient of variation (CV) for each genus was calculated. A small constant (1 × 10 ¹) was added to the denominator to avoid division-by-zero errors in cases of extremely low abundance values. Higher CV values were interpreted as indicating greater inter-individual variability and, therefore, lower microbial stability.

Differences in core stability were assessed across groups and time points using the nonparametric Wilcoxon rank-sum test. Resulting *P*-values were adjusted for multiple comparisons with F to evaluate statistical significance.

To further characterise the most stable core taxa, the top five genera with the lowest CV values at baseline (D0) were identified for each group. The longitudinal variation of these genera was tracked across time points and compared with the mean CV across all core genera.

#### Assessing the individuality of the microbial communities

To assess temporal correlation in microbial community structure across time points for a particular animal, we first calculated beta diversity using the Bray–Curtis dissimilarity metric on absolute abundance data from faecal samples separately (*phyloseq::distance()* function). The resulting distance matrix was converted into a data frame and enriched with metadata, including animal ID and sampling day. We reshaped the data into long format using the *melt()* function to facilitate mixed-effects modelling. Then, we computed Spearman’s ρ values for each animal at consecutive time points, ensuring comparisons were restricted to individuals sampled at both time points. This yielded a series of pairwise correlation values reflecting microbial similarity over time, summarised in a time-resolved correlation profile. Higher correlation values indicated more stable microbial communities, while lower values suggested greater variability.

#### Dynamic microbial dispersion and divergence between individuals

We first used the *vegdist()* function from the vegan package to calculate inter-individual variation over time for faecal microbiota, using pairwise Bray-Curtis dissimilarities. Specifically, we focused on the inter-individual variation by calculating the mean Bray-Curtis distance between all pairs of individuals at each time point. This allowed us to quantify differences in microbial community composition across individuals over time. Next, we used the *mandist()* function from the vegan R package to compute the mean distances.

We also used the similarity analysis (ANOSIM) to test for intragroup dispersion. ANOSIM is a permutation-based test in which the null hypothesis is that within-group distances are not significantly smaller than between-group distances. The test statistic (R) ranges from -1 to 1, with 1 indicating that all samples within a group are more similar to one another than to any representative of another group. R is ≈ approximately 0 when the null hypothesis is true, meaning that the distances within and between groups are the same on average. Because multiple comparison corrections for ANOSIM were unavailable, the number of permutations used in those calculations increased to 9,999.

#### Resistance of the microbial communities

The ability to remain essentially unchanged despite disturbances is referred to as resistance [101]. In this study, microbial ecosystem resistance was quantified to assess the temporal stability of the faecal microbiota across groups. Resistance was calculated from presence/absence-transformed ASV data derived from absolute-abundance phyloseq objects. ASV tables were extracted and converted to binary format, where taxa were considered present if their abundance exceeded zero. Sample metadata, specifically the Time variable, was used to group samples by time. Resistance at a given time point *t* was defined as one minus the mean Jaccard dissimilarity between samples collected at time *t* and those collected at the subsequent time point *t+1*. This approach captures the proportion of shared taxa between adjacent time points, with higher resistance values indicating greater community stability. To compute this, samples from each pair of consecutive time points were subset, and their ASV matrices were merged. A Jaccard dissimilarity matrix was then calculated using the *vegdist()* function from the vegan package. From this matrix, pairwise distances between samples from *t* and *t+1* were extracted, and the mean dissimilarity was computed. Resistance was then calculated as one minus this mean value and assigned to the time point *t*. This approach is based on the established ecological theory of community similarity [102].

#### Microbial Differential taxonomic abundance analysis (MaAsLin2)

Differential abundance analysis was used to identify taxonomic changes produced in the microbiota over the duration of the study. To achieve this, Multivariate Association with Linear Models analysis (MaAslin2) [103] (v.1.15.1) was performed. To evaluate diet-related effects independently of temporal structure, analyses were conducted separately for each sampling time point (D0–D32).

The Control group was used as the reference category. Relative abundances were normalised using total sum scaling (TSS) and subsequently log-transformed. To reduce noise and limit the influence of rare taxa, features with a minimum relative abundance below 1×10 or present in fewer than 5% of samples were removed. Statistical significance was assessed using linear models, and p-values were corrected for multiple testing using the false discovery rate (FDR). Associations with an FDR *q*-value < 0.05 were considered significant. This procedure was applied independently for each time point and experimental group.

Significant associations were then aggregated into a merged dataset. Taxonomic annotations at the Order, Family, and Genus levels were retained (Order_Family_Genera). Taxa lacking classification or flagged as NA were labelled as *Unclassified_ASV*. The direction of each association was determined using the regression coefficients: positive coefficients indicated higher abundance in the Chicory group, whereas negative coefficients indicated higher abundance in the Control group.

#### Analysis of Environmental Drivers (Envfit) on microbiota structure

Fits an Environmental Vector or Factor onto an Ordination (Envfit) was performed to identify the variables that significantly explained the variation in the microbial community structure. This analysis was performed using the *envfit()* function from the vegan package [104] (v2.7-2) in R.

Environmental variables from the phyloseq object used in the analysis were selected for their biological significance. Short-chain fatty acids were decomposed into their individual components to provide a more detailed description (Acetate, Propionate, Isobutyrate, Butyrate, Isovalerate, and Valerate, respectively). The ordination was calculated from the microbiome’s beta-diversity. Bray-Curtis dissimilarity matrices and PCoA from the β-diversity section were reused. The *envfit* function was applied to the PCoA vectors to project environmental variables onto the ordination space. Permutation tests (9999 permutations) were conducted to assess the statistical significance (*P* < 0.05) of each environmental variable. The analysis was conducted independently for each time point across the study period. Drivers were mapped on the ordination plot as vectors. The length, thickness, and direction of each variable represent the magnitude and orientation of its correlation with the bacterial community composition. Significant variables were plotted, and their R^2^ values were used to assess the magnitude of the association with modulation of microbial composition.

#### qPCR Quantification of Bacterial biomass in faeces samples

As detailed in Mach et al. (2025)[105], bacterial concentrations in faecal samples were quantified by qPCR on a QuantStudio 12K Flex platform (Thermo Fisher Scientific, Waltham, United States). Primers targeting total bacteria (forward: 5′-CAGCMGCCGCGGTAANWC-3′; reverse: 5′-CCGTCAATTCMTTTRAGTTT-3′) were obtained from Eurofins Genomics (Ebersberg, Germany). PCR products generated from the corresponding amplicons were used to prepare a seven-point, ten-fold serial dilution to construct the standard curve. For bacterial quantification, the dilution series ranged from 2.25 × 10□to 2.25 × 10¹³ copies per µg of DNA. Quantitative PCR assays were carried out in 20□µL reaction volumes, each containing 10□µL of Power SYBR Green PCR Master Mix (Thermo Fisher), 2□µL of either template DNA or standard (0.5□ng/µL), and 0.6□µM of each primer, yielding a final primer concentration of 200□mM for bacterial assays. Thermal cycling consisted of an initial denaturation at 95□°C for 10□min, followed by 40 cycles of 95□°C for 15□s and 60□°C for 60□s for combined annealing and extension. A melt-curve analysis was performed at the end of each run by gradually increasing the temperature from 60□°C to 95□°C to verify amplification specificity. All reactions were run in triplicate, and bacterial copy numbers in faecal samples were calculated using a standard curve generated from purified amplicons.

Copy numbers were estimated using the standard formula based on nucleotide molecular mass and amplicon length: copies per nanogram = (NL × A × 10□□) / (n × mw), where NL is Avogadro’s constant (6.02 × 10²³ molecules/mol), A is the DNA mass (ng), n is the amplicon size (bp), and mw is the average molecular weight per base pair.

Bacterial abundance (biomass) was log□□-transformed to meet the assumptions of parametric analyses, specifically normality and homoscedasticity of residuals. Results are therefore presented as log□□gene copies per gram of wet faeces (Supplementary Table S1). Comparisons of biomass across sampling sites were performed using the Kruskal–Wallis test, implemented in the *kruskal.test()* function of the stats package in R.

#### Cyathostomin larval DNA extraction from faecal samples

Based on larval counts, a volume corresponding to 1,000 larvae was sampled from each animal and time point. However, 10 samples contained fewer than 1,000 larvae in 2 mL. Tubes with larvae were subsequently centrifuged at 10,000 × g for 2 minutes to pellet the larvae, and the supernatant was removed. Larvae DNA extraction was performed using the NucleoMag Tissue kit (Macherey-Nagel). For each sample, three 3 mm steel grinding beads, 200 µL of lysis buffer T1, and 25 µL of proteinase K were added. Samples were homogenised using a TissueLyzer bead beater (Retsch) for 2 minutes at 20 Hz, then incubated at 56°C for 1 hour with shaking at 600 rpm. DNA purification was then carried out according to the manufacturer’s protocol using the KingFisher Apex automated system (Thermo Fisher Scientific), with elution in 100 µL of MB6 buffer.

DNA concentrations were measured using the Quantifluor reagent (Promega) on a FluoStar Omega spectrofluorometer (BMG Labtech). Extracts were subsequently normalised, where possible, to 5 ng/µL in a final volume of 40 µL (average concentration: 4.1 ng/µL across all samples) using a Biomek i7 automated workstation. The KingFisher Apex and Biomek i7 instruments were operated at the Symbiotron platform (University of Lyon 1, France).

#### ITS-2 amplification, sequencing and nemabiome analysis

DNA was extracted from cultured third-stage NGI larvae (L3). Of the 154 initial samples, 149 provided sufficient larval material for nemabiome analysis. Whenever possible, an aliquot corresponding to approximately 1,000 larvae was recovered from each sample. Larvae were pelleted by centrifugation, and genomic DNA was extracted using the NucleoMag Tissue kit (Macherey-Nagel) following mechanical disruption with a TissueLyser (Retsch) and proteinase K digestion. DNA purification was performed on a KingFisher Apex automated workstation (Thermo Fisher Scientific), and DNA was eluted in 100 µL of MB6 buffer. DNA concentrations were quantified using the Quantifluor dsDNA assay (Promega) on a FLUOstar Omega fluorometer (BMG Labtech), and extracts were normalised to 5 ng µL□¹ whenever possible using a Biomek i7 automated liquid-handling workstation (Beckman Coulter).

The ITS-2 region of ribosomal DNA was amplified using modified NC1 and NC2 primers that target the conserved 5.8S and 28S rDNA regions. Primers incorporated Illumina adapter sequences together with 0-3 random nucleotides to increase sequence diversity during cluster generation. Equal mixtures of four forward and four reverse primer variants were used for PCR amplification with KAPA HiFi HotStart DNA polymerase. PCR amplification, library preparation, and sequencing were performed by IGA Technology Services (Udine, Italy) using a custom amplicon sequencing workflow. Libraries were sequenced on an Element AVITI™ platform using paired-end 2 × 300 bp chemistry (Custom Amplicon-Seq, AVITI PE300), targeting approximately 50,000 paired-end reads per sample.

Sequence data were processed following a DADA2-based nemabiome pipeline adapted from elsewhere [106]. Primers were removed using Cutadapt prior to quality filtering and denoising in DADA2. Reads were filtered using the filterAndTrim() function with a minimum length of 50 bp and maximum expected error thresholds of 2 and 5 for forward and reverse reads, respectively. Amplicon sequence variants (ASVs) were inferred using the dada() algorithm, paired-end reads were merged, and chimeric sequences were removed using removeBimeraDenovo(). Taxonomic assignment was performed using the nematode ITS-2 reference database (https://www.nemabiome.ca/), and complementary classification approaches. ASVs were first classified using the IDTaxa machine-learning classifier with a confidence threshold of 60. Low-confidence assignments were re-evaluated using a second IDTaxa pass (threshold = 30), followed by Bayesian classification using assignTaxonomy() in DADA2. The resulting ASV table was curated by removing singletons; ASVs detected exclusively in negative controls; putative cross-contamination or tag-jumping artefacts; ASVs with fewer than 10 reads; and samples with fewer than 1,000 reads after filtering. One sample was excluded during this step. The final curated ASV table was used for all downstream analyses.

#### Bacteriome – Nemabiome Association Analysis

Global associations between shifts in bacterial and nematode taxa were evaluated using distance-based correlation approaches. Bacterial communities were analysed at the genus level, while nematode communities were analysed at the species level.

Taxonomic agglomeration was performed at each taxonomic category. Only samples shared between phyloseq objects were retained, and the phyloseq components were reordered to maintain the same sample order. Community dissimilarity matrices were calculated for the dataset using Bray-Curtis distances. Concordances between community structures were assessed using the Mantel test with Spearman correlations (999 permutations). The strength of the association was quantified using both correlation coefficients.

Deeper structural congruence between ordination spaces was evaluated using Procrustes analysis. Principal coordinate analysis was performed on Bray-Curtis distance matrices using classical multidimensional scaling. Procrustes statistics (M^2^) and permutation significance values were calculated with 999 permutations.

Sparse correlations for Compositional data (SparCC) were used to analyse the statistical significance of these potential associations. Core taxa were targeted to reduce the sparse, low-abundance features and focus on analysing the highly abundant and prevalent members of the community. Pairwise correlations were calculated using the SparCC algorithm. To assess the statistical significance, a model was performed using 999 permutations. Empirical p-values were estimated and adjusted with FRD correction.

Association patterns were visualised with heatmaps, with positive and negative correlations represented by a colour-coded gradient.

## Data Availability declaration

16S rRNA gene sequences are available in the DDBJ/EMBL/GenBank under the BioProject ID PRJNA1472508, titled “Chicory Reshapes the Equine Holobiont**”** at the NCBI Sequence Read Archive (SRA) (www.ncbi.nlm.nih.gov/sra/PRJNA1472508). The BioSamples SAMN60498732 to SAMN60498886 correspond to the 156 individual faecal samples collected for 16S rRNA amplicon sequencing within the BioProject (PRJNA1472508). Each BioSample name encodes the animal identifier, the treatment group, and the sampling time point across the longitudinal design. Together, these samples represent the full temporal series of microbiota profiles generated from naturally parasitised yearling and two-year-old horses grazing pastures with contrasting chicory levels. The metadata associated with each BioSample includes the horse ID, age class, treatment allocation, sampling date, and experimental phase, ensuring complete traceability of every sample throughout the study. Temporary Submission ID: SUB16217194.

The ITS-2 amplicon sequencing data generated to characterise the equine gastrointestinal nemabiome are available in the DDBJ/EMBL/GenBank databases under BioProject accession PRJNA1482472, entitled “The Equine Nemabiome”, at the NCBI SRA (https://www.ncbi.nlm.nih.gov/sra/PRJNA1482472).

The BioSamples SAMN61125566 **to** SAMN61125714 correspond to the 149 individual faecal samples collected for ITS2 amplicon sequencing. Each BioSample name encodes the individual horse identifier, and the associated metadata include the horse ID, treatment group, sampling time point, and sequencing library, ensuring complete traceability of each sample throughout the study.

The complete metadata associated with this project, together with the phyloseq objects for the microbiota and nemabiome datasets and all analysis scripts, are available in the INRAE institutional data repository powered by Dataverse and are publicly available as of the date of publication, with DOI: https://doi.org/10.57745/OQFNPL.

The lead contact can provide any additional information required to reanalyse the data reported in this paper upon request.

## Acknowledgments

The authors thank Hortensia Robert, Agathe Garcia, and Soledad for their help with the fieldwork and larval selection, and Guenaelle Derolez for her help with the qPCR. We thank the MIGALE bioinformatics facility for providing support, computing, and storage resources (https://doi.org/10.15454/1.5572390655343293E12; Jouy-en-Josas, France).

## Funding Declaration

This study was supported by the *Institut Français du Cheval et de l’équitation* (IFCE), whose funding contributed to laboratory analyses and field experimentation, and by Carnot France Future Élevage, which contributed to sequencing and cytokine and biochemical analyses.

## Author Contribution

NM wrote the main manuscript and performed the statistical analyses of the host responses. SM generated the main figures and conducted the statistical analyses for the microbiome and nemabiome datasets. JM contributed to the study design and participated in the experimental work. JAR was responsible for the Chamberet facilities where the experiment was conducted and contributed to the experimental design and work. DB and MASB participated in the experimental work. DB performed the microbiota bioinformatics analysis. GP and MMI assisted with SCFA analyses and sample preparation. AV and LL, as experts in animal behaviour, advised on and refined the behavioural observation protocol. ARW provided expertise in immunology, and ER analysed the biochemical parameters and Th2 immune responses. GB, an expert in parasitology, HH, OC and CR, experts in molecular biology and CB and GY, experts in nemabiome analyses, were jointly responsible for generating the FEC, larvae, and nemabiome datasets. GF and NM designed the experiment and secured the project funding. GF also conducted the botanical component of the study. All authors reviewed and approved the final manuscript.

## Competing Interest declaration

The authors declare that they have no competing interests.

## ADDITIONAL FIGURES

**Additional Figure 1.**
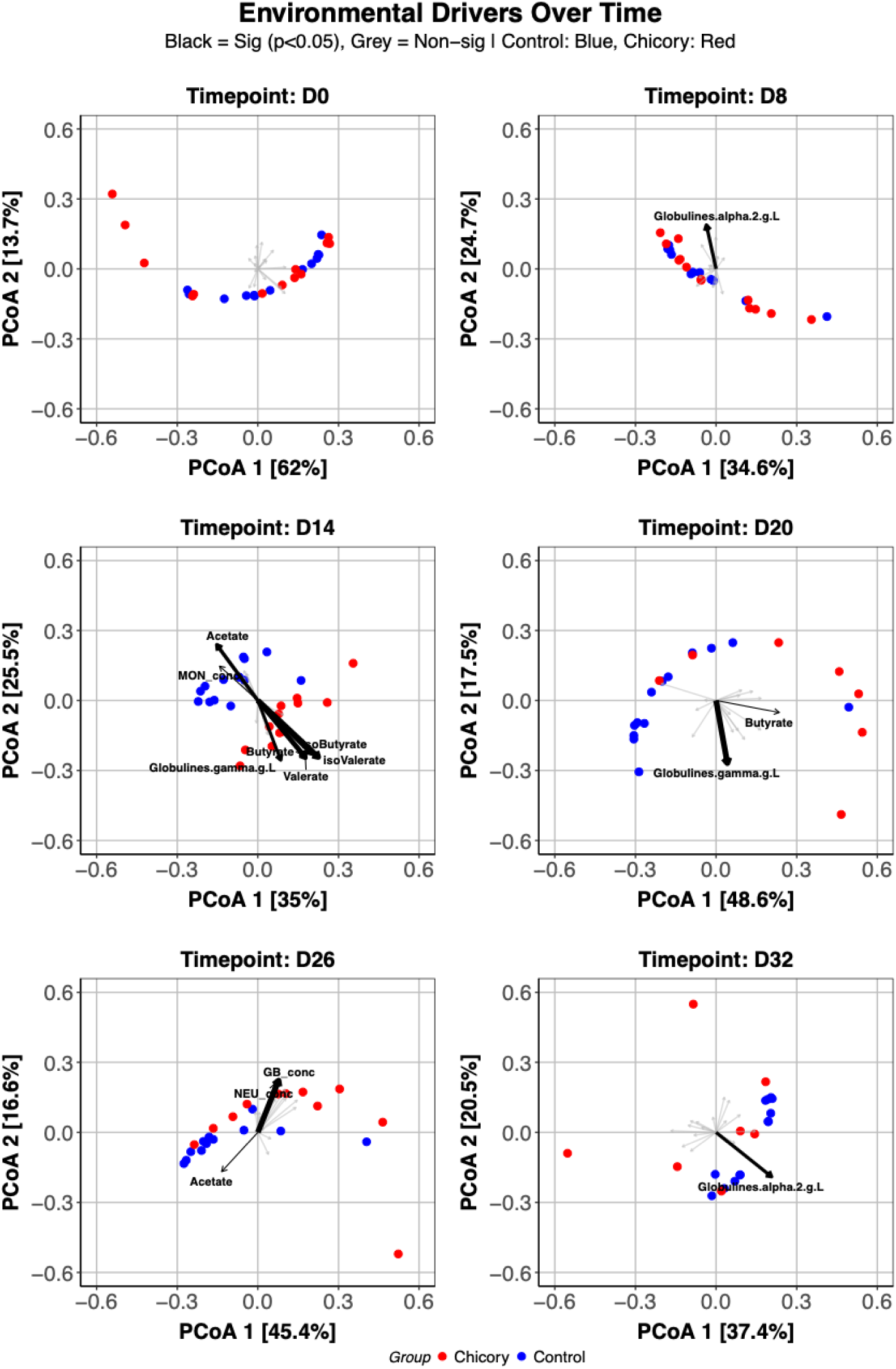
Environmental drivers of microbiota structure across the grazing trial. Ordination plots (PCoA based on Bray–Curtis dissimilarities) showing the temporal evolution of microbial community structure in control (blue) and chicory-fed (red) horses from Day□0 to Day□32. At each time point, vectors represent the environmental and host variables significantly associated with community composition according to the *envfit* model (*P*□<□0.05; variables in black), while non-significant variables are shown in grey.

**Additional Figure 2.**
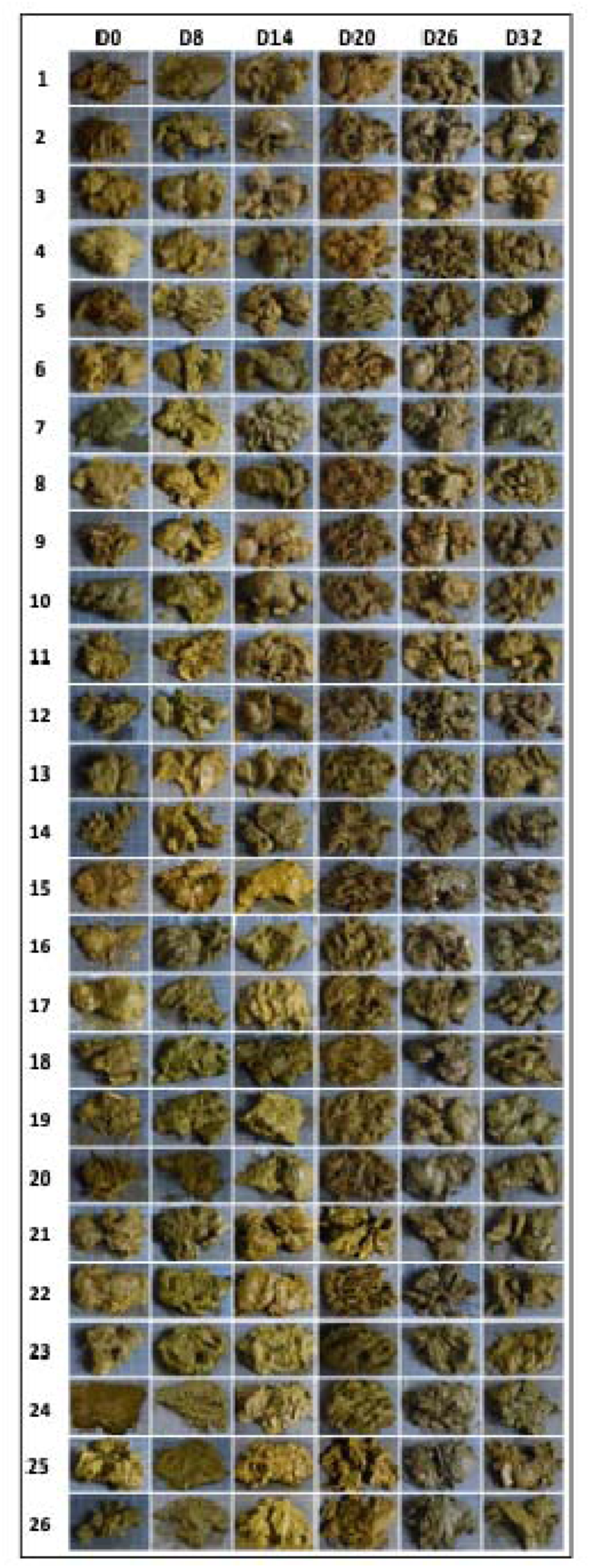
Macroscopic characteristics of faeces collected from control and chicory-fed animals over time. Samples are numbered from 1 to 26: samples 1–13 correspond to control animals, while samples 14–26 correspond to chicory-fed animals. Representative images illustrate changes in fecal appearance across the different time points (D0, D8, D14, D20, D26, and D32).

## ADDITIONAL TABLES

**Additional Table 1.** Experiment metadata. Individual-level zootechnical, physiological, haematological, biochemical, immunological, and faecal fermentation parameters were measured in horses throughout the experimental period. For each horse, the table reports identification data, sampling information, and core physiological indicators.

**Additional Table 2.** Amplicon sequence variants (ASVs) of the nemabiome with taxonomic annotation up to species level and corresponding raw read counts. This table lists all detected nemabiome ASVs, providing their taxonomic classification from higher ranks down to species level where possible, along with the associated raw sequence counts per sample. Raw counts represent the unnormalized abundance of each ASV and are provided to allow transparency and reproducibility of downstream analyses, including diversity estimation and community structure comparisons.

**Additional Table 3.** Daily inter-group structural differentiation of the nemabiome assessed using Bray–Curtis ANOSIM (Analysis of Similarities). For each sampling day, the table reports the ANOSIM R statistic, p-value, number of permutations (n = 9,999), FDR-adjusted p-value, and the resulting significance level. The R statistic reflects the degree of separation between experimental groups: values close to 1 indicate strong between-group dissimilarity, values near 0 indicate no separation, and negative values indicate greater within-group than between-group variation.

**Additional Table 4.** Differentially abundant nematode taxa (MaAsLin2 analysis) between chicory and control groups at each time point. Results from MaAsLin2 analysis identifying nematode taxa significantly associated with chicory versus control across sampling times (D8, D20, D26, D32). The table includes taxonomic classification (order, family, genus, species), model coefficients (coef), standard errors (stderr), raw *P*-values, and false discovery rate–adjusted q-values (qval). Negative coefficients indicate taxa enriched in the control group, whereas positive coefficients indicate enrichment in the chicory group. Direction denotes the group in which each taxon is significantly more abundant.

**Additional Table 5.** Bacterial amplicon sequence variants (ASVs) from 16S rRNA gene sequencing with taxonomic annotation up to species level and corresponding abundance counts. This table presents all bacterial ASVs identified through 16S rRNA sequencing, including their taxonomic classification from higher ranks to species level when available, along with the associated abundance sequence counts per sample. Abundance counts represent the normalised abundance of each ASV.

**Additional Table 6.** Relative abundance and prevalence of microbial genera across treatments and timepoints with ecological classification. This table presents the mean relative abundance and prevalence of microbial genera across experimental treatments and timepoints. Each row corresponds to a genus (or the lowest available taxonomic assignment) within a specific treatment group and sampling time. Genus indicates the taxonomic classification (family_genus or unclassified group). Mean_Pct represents the average relative abundance (%) of each genus across samples, while SD_Pct reflects the associated standard deviation, indicating variability. Prevalence corresponds to the proportion of samples in which a given genus was detected (range: 0–1). Treatment specifies the experimental group (e.g., Control or dietary intervention), and Timepoint indicates the sampling time (e.g., D0 for baseline and D32 for post-treatment). Genera are classified into ecological categories: Core taxa are highly prevalent and consistently detected across samples; HPA (High Prevalence Abundant) taxa are frequently detected with moderate abundance; LPA (Low Prevalence Abundant) taxa are rare and/or present at low abundance.

**Additional Table 7.** Differentially abundant bacterial taxa (MaAsLin analysis) between chicory and control groups at each time point. Results from MaAsLin analysis showing bacterial taxa significantly associated with chicory versus control across sampling times (D8, D14, D20, D26, D32). The table includes taxonomic classification (order, family, genus), model coefficients (coef), standard errors (stderr), raw *P*-values, and false discovery rate–adjusted q-values. Positive coefficients indicate taxa enriched in the chicory group, whereas negative coefficients indicate enrichment in the control group. Direction indicates the group in which each taxon is significantly more abundant.

**Additional Table 8.** SparCC correlation analysis between bacterial taxa (16S rRNA sequencing) and nematode taxa across experimental groups. This table presents pairwise correlations between bacterial taxa identified by 16S rRNA gene sequencing and nematode taxa from the nemabiome, calculated using the SparCC method. For each bacterium–nematode pair, the table reports the SparCC correlation coefficient (indicating the strength and direction of the association), the FDR-adjusted p-value, and the experimental group (e.g., Control, Chicory). Positive coefficients indicate co-occurrence, whereas negative coefficients suggest inverse relationships. Adjusted *P*-values account for multiple testing, allowing identification of statistically robust associations.

**Additional Table 9.** Sequences of primers, probes, and concentrations used for the quantification of cytokines and housekeeping genes. IL: interleukin; GAPDH: glyceraldehyde-3-phosphate dehydrogenase; ACTB: actin β; F: forward primer; R: reverse primer; P: probe.

## Notes

https://doi.org/10.57745/OQFNPL .

### Competing Interest Statement

The authors have declared no competing interest.

